# Synthetic regulatory genomics uncovers enhancer context dependence at the *Sox2* locus

**DOI:** 10.1101/2022.06.20.495832

**Authors:** Ran Brosh, Camila Coelho, André M. Ribeiro-dos-Santos, Gwen Ellis, Megan S. Hogan, Hannah J. Ashe, Nicolette Somogyi, Raquel Ordoñez, Raven D. Luther, Emily Huang, Jef D. Boeke, Matthew T. Maurano

## Abstract

Expression of *Sox2* in mouse embryonic stem cells (mESCs) depends on a distal regulatory cluster of DNase I hypersensitive sites (DHSs), but their individual contributions and degree of independence remain a mystery. Here, we comprehensively analyze the regulatory architecture of *Sox2* at its endogenous locus using Big-IN to scarlessly integrate DNA payloads ranging up to 143 kb. We analyzed 83 payloads incorporating deletions, rearrangements, and inversions affecting single or multiple DHSs, as well as surgical alterations to transcription factor (TF) recognition sequences. Multiple mESC clones were derived for each payload, sequence-verified, and analyzed to establish the necessity and sufficiency of genomic features for *Sox2* expression. We found that two LCR DHSs comprising a handful of key TF recognition sequences were each sufficient to autonomously sustain significant expression in mESCs. However, three additional LCR DHSs were entirely context-dependent, in that they showed no activity alone but could dramatically augment activity of the core DHSs. Our synthetic regulatory genomics approach demonstrates that composite regulatory elements can be reduced to a tractable set of essential sequence features, and is readily scalable to investigate regulatory architecture at other key loci genome-wide.

## Introduction

Many developmentally important genes lie near clusters of highly cell-type specific DNase I hypersensitive sites (DHSs), typified by the β-globin locus control region (LCR)^1–3^. More recently, ‘super-enhancers’ have been proposed to exhibit exceptionally cooperative binding, acting as developmental switches that regulate key TFs^4–6^. Exploration of the broad assumptions involved has been impeded by the challenges of large-scale genomic engineering, leaving fundamental questions unanswered. Individual constituent DHSs demonstrate enhancer activity in transient expression assays^4, 7^, but it is less clear whether they act independently or synergistically in composite elements^8–11^. Furthermore, nearby enhancers may provide redundancy rather than augmenting gene expression per se^12–15^. Therefore, there is a need for new approaches to investigate the architecture of complex regulatory elements.

*Sox2* encodes a key TF that regulates mouse embryonic stem cell (mESC) self-renewal and pluripotency^16^. The *Sox2* gene is surrounded by a proximal enhancer cluster, but its expression in mESCs relies on an LCR comprising multiple DHSs located 100 kb downstream^17, 18^. The effect of the LCR on *Sox2* expression is influenced by distance and/or intervening CTCF sites^19–21^. Surgical inversion of a single CTCF recognition sequence within the *Sox2* LCR affects chromatin architecture but not *Sox2* expression in mESCs^22^, suggesting that proximity and expression might be separate functions^23, 24^. The *Sox2* locus therefore provides a natural model for dissecting how the composition and configuration of complex regulatory elements determine transcriptional activity. However, previous studies have focused on transient reporter plasmids which may not recapitulate function at the endogenous locus, and simple deletion analyses which do not address sufficiency or interaction of individual elements.

We have recently developed the Big-IN platform to enable scarless genome rewriting with payloads exceeding 100 kb^25^. Big-IN includes two engineering steps: first, CRISPR/Cas9 is used to target and replace an allele of interest with a landing pad (LP) (**Fig. 1a**). Second, the LP is replaced by a transfected DNA payload using recombinase-mediated cassette exchange (RMCE), and payload-harboring cells are isolated using a positive/negative selection strategy. A comprehensive sequencing verification pipeline accompanies each step to ensure on-target single copy integration and lack of unexpected changes (**Fig. 1b**).

**Fig. 1.**
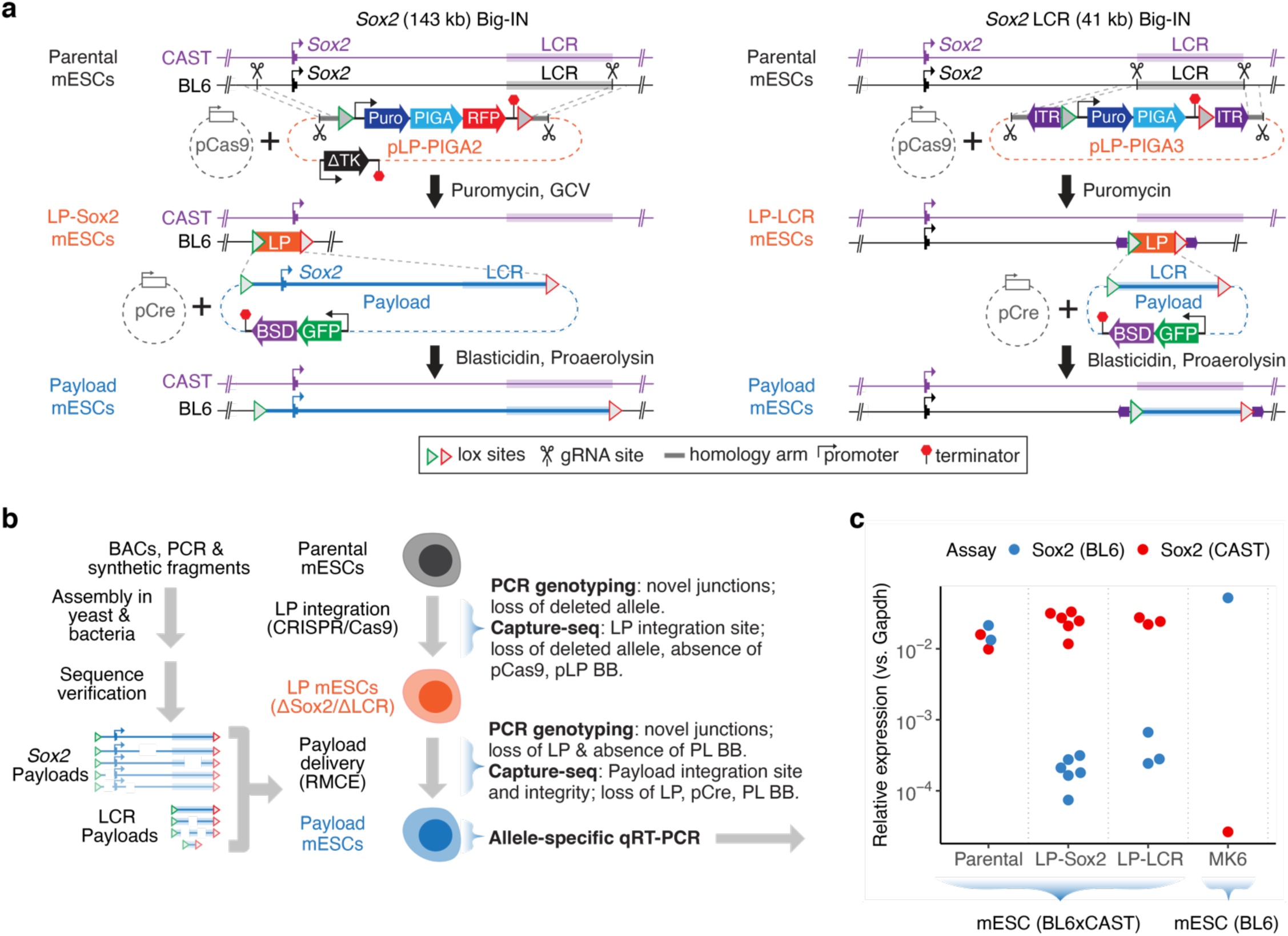
A synthetic regulatory genomics pipeline for investigation of *Sox2* locus architecture. **a.** mESC engineering strategy. The BL6 allele of the 143-kb *Sox2* locus (left) or the 41-kb *Sox2* LCR (right) were replaced with LP-PIGA2 or LP-PIGA3, respectively. LP integration was aided by Cas9 using a pair of gRNAs targeting both the replaced allele and the LP plasmid and by short homology arms that facilitate homology-directed repair. LP mESCs were selected with puromycin, while ganciclovir (GCV) selects against integration of the LP backbone (BB). Cre recombinase-mediated cassette exchange (RMCE) enabled replacement of each LP with a series of payloads. Transfected cells were transiently selected with blasticidin, followed by counterselection of LP mESCs cells with proaerolysin. **b.** Schematic of DNA assembly, mESC engineering, verification, and analysis pipelines. **c.** Allele-specific qRT-PCR assay for *Sox2* expression. BL6 and Castaneus (CAST) expression was measured using allele-specific primers in parental BL6xCAST mESCs, in mESCs with deletion of the 143-kb *Sox2* locus (ΔSox2) or 41-kb deletion of the LCR (ΔLCR), as well as in BL6 MK6 mESCs.

Here we use Big-IN to rewrite the mouse *Sox2* locus with designer payloads incorporating deletions, inversions, translocations, and mutations of single and multiple DHSs. We show that the LCR is a complex element whose constituent DHSs are only partially interchangeable and exhibit unexpected dependencies not previously predicted by reporter assays or deletion analyses. We show that the contribution of individual TF activity depends critically on their local context. Future application of this synthetic regulatory genomics approach to assess regulatory architecture promises to cast new light on the function of key loci genome-wide.

## Results

### Synthetic regulatory genomics of the *Sox2* locus

To enable interrogation of regulatory element function at the murine *Sox2* locus, we integrated Big-IN LPs into the BL6 allele of C57BL6/6J × CAST/EiJ (BL6xCAST) F1 hybrid mESCs. We targeted independent LPs to replace both the 143-kb *Sox2* locus (LP-Sox2) and the 41-kb region surrounding the *Sox2* LCR (LP-LCR) (**Fig. 1a**). We performed comprehensive quality control (QC) using PCR genotyping and sequencing, which identified clones with single-copy, on-target LP integration (**Supplementary Fig. S1**, and **Table 1**). Using real-time quantitative reverse transcription PCR (qRT-PCR) with allele-specific primers, we confirmed biallelic *Sox2* expression in parental cells, ablation of BL6 *Sox2* expression in LP-Sox2 mESCs, and near total loss of BL6 *Sox2* expression in LP-LCR mESCs, recapitulating previous reports (**Fig. 1c**)^17, 18^. Total loss of the BL6 *Sox2* allele was well-tolerated, with no discernable impact on cell morphology or growth rate.

**Table 1.**
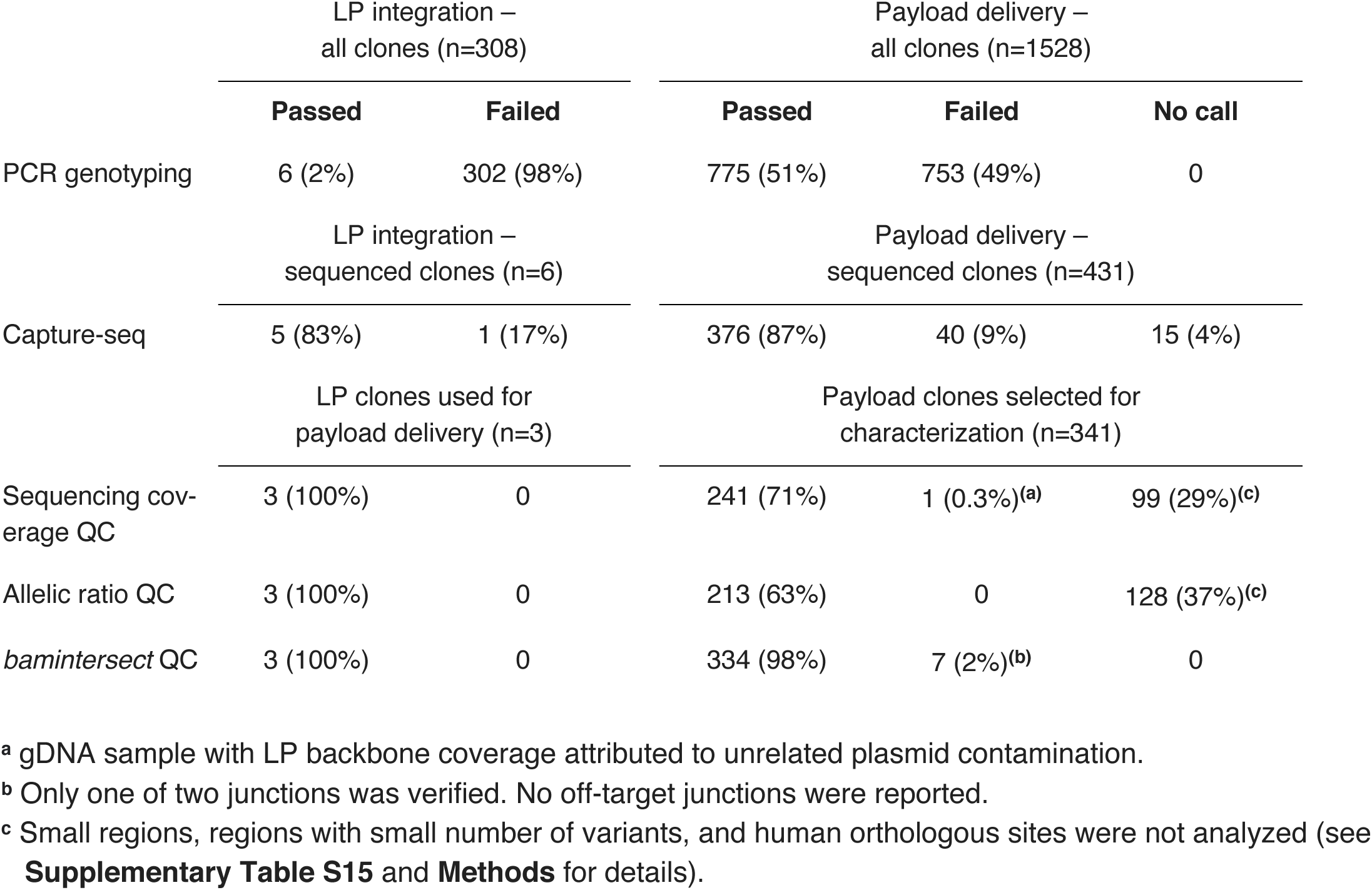
Engineered mESC clones quality control summary. Landing pad (LP) and payload mESC clones were screened first using PCR genotyping and then Capture-seq. Clones included in this study and profiled for *Sox2* expression were further verified by computational analyses of sequencing coverage depth and allelic ratios (**Supplementary** Table S15) and *bamintersect* integration site analysis (**Supplementary** Table S17). Percentages are relative to the number of clones at each stage.

|  | LP integration –<br>all clones (n=308) |  | Payload delivery –<br>all clones (n=1528) |  |  |
| --- | --- | --- | --- | --- | --- |
|  | Passed | Failed | Passed | Failed | No call |
| PCR genotyping | 6 (2%) | 302 (98%) | 775 (51%) | 753 (49%) | 0 |
|  | LP integration –<br>sequenced clones (n=6) |  | Payload delivery –<br>sequenced clones (n=431) |  |  |
|  | Passed | Failed | Passed | Failed | No call |
| Capture-seq | 5 (83%) | 1 (17%) | 376 (87%) | 40 (9%) | 15 (4%) |
|  | LP clones used for<br>payload delivery (n=3) |  | Payload clones selected for<br>characterization (n=341) |  |  |
|  | Passed | Failed | Passed | Failed | No call |
| Sequencing coverage QC | 3 (100%) | 0 | 241 (71%) | 1 (0.3%)(<sup>a</sup>) | 99 (29%)(<sup>c</sup>) |
| Allelic ratio QC | 3 (100%) | 0 | 213 (63%) | 0 | 128 (37%)(<sup>c</sup>) |
| *bamintersect* QC | 3 (100%) | 0 | 334 (98%) | 7 (2%)(<sup>b</sup>) | 0 |

<sup>a</sup> gDNA sample with LP backbone coverage attributed to unrelated plasmid contamination.
<sup>b</sup> Only one of two junctions was verified. No off-target junctions were reported.
<sup>c</sup> Small regions, regions with small number of variants, and human orthologous sites were not analyzed (see **Supplementary Table S15** and **Methods** for details).

We assembled DNA payloads for Big-IN delivery (size = 177 to 142,667 bp) from a BL6 mouse BAC (bacterial artificial chromosome) encompassing the full *Sox2* locus or from chemically synthesized DNA (**Supplementary Table S1**). We verified the resulting payload BACs through sequencing (**Supplementary Table S2**), delivered them to LP mESCs, isolated multiple mESC clones for each payload, and comprehensively verified single-copy targeted payload integration using targeted capture sequencing (Capture-seq) (**Table 1, Supplementary Table S3**). Approximately 50% of isolated delivery clones passed PCR genotyping, and the majority of those (∼87%) passed subsequent Capture-seq screening. All 341 mESC clones characterized in this study were further confirmed by systematic analyses of sequence coverage depth, allelic ratios, and integration site (**Table 1**). The high proportion of on-target payload delivery attests to the fidelity of RMCE and the efficiency of the positive and negative selection for Big-IN delivery.

To dissect the architecture of the *Sox2* locus, we delineated 28 distinct regulatory regions (**Fig. 2a**) through inspection of genomic annotations including DNA accessibility and TF occupancy data in mESC. We delivered payloads including single and multiple deletions of these regions to LP-Sox2 mESCs. We measured *Sox2* expression from the engineered allele relative to the unedited CAST allele as an internal control (**Supplementary Fig. S2**). Delivery of the wild-type (*WT*) locus fully rescued *Sox2* expression, whereas payloads lacking the LCR (*ΔLCR*) showed minimal expression (**Fig. 2b**, **Supplementary Table S4**). Single deletion of DHS7 or DHS15 or the combined deletion of DHSs 10-11 have been reported to show no expression difference^17^. Yet we found that deletion of DHSs 10-16 downstream of *Sox2* led to a 22% reduction in expression. A larger deletion of the entire 81-kb intervening region between *Sox2* and its LCR (including DHSs 10-18) showed expression comparable to WT, possibly as the reduced distance between *Sox2* and its LCR partially compensates for loss of proximal regulatory elements as reported previously^20^. Deletion of the proximal region upstream of *Sox2* (DHSs 1-8) also reduced expression. Combined deletion of both the upstream and downstream proximal DHSs led to even lower *Sox2* expression than either single deletion. A more restricted deletion of DHSs 1-8 with just DHSs 10 & 15 showed a similar reduction in activity, while a combined deletion of DHSs 1-8 with CTCF17 alone or with CTCF sites 13-14 showed little effect. We conclude that the effect of the downstream deletion is largely attributable to DHSs 10 & 15, and note that both of which were found active in STARR-seq and luciferase reporter assays^17, 26^ (**Fig. 2a**). Our results suggest that some of the *Sox2*-proximal DHSs are redundant and show no effect unless deleted together.

**Fig. 2.**
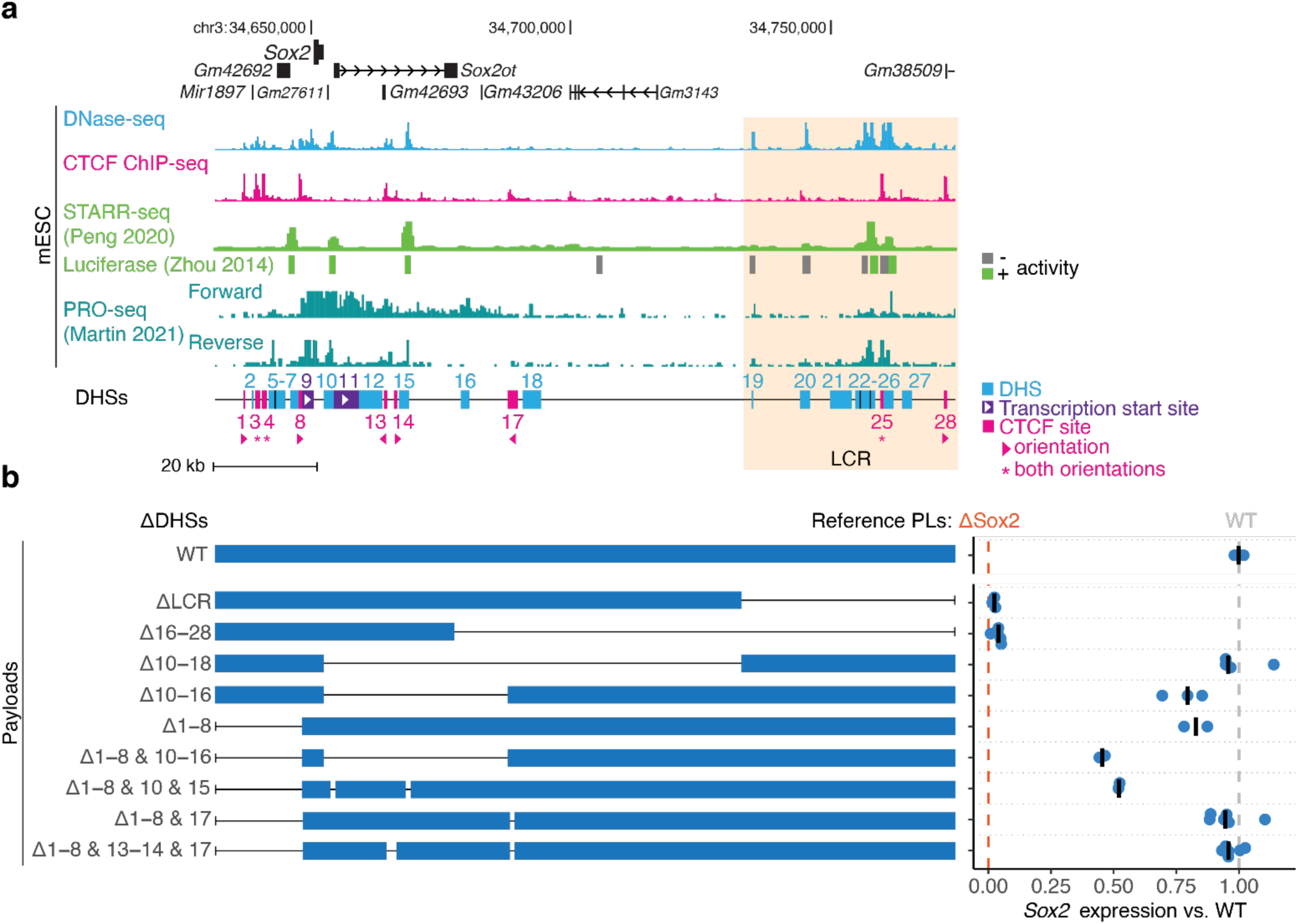
Redundancy of proximal enhancers at the *Sox2* locus. **a.** Schematic of the *Sox2* locus. Shown are DNase-seq, CTCF ChIP-seq, reporter assay (STARR-seq^26^ and luciferase^17^), and PRO-seq^31^ data from mESCs. Orientation of CTCF recognition sequences is indicated where applicable. **b.** *Sox2* expression analysis for payloads delivered to the *Sox2* locus in LP-Sox2 mESCs. Blue rectangles demarcate genomic regions included in each payload. Each point represents the expression of the engineered *Sox2* allele in an independent mESC clone. Bars indicate median. Expression was scaled between 0 (*ΔSox2*) to 1 (*WT LCR*).

### Dissection of the *Sox2* LCR

We next focused on a 41-kb region surrounding the *Sox2* LCR containing a total of 10 DHSs and CTCF sites (**Fig. 3a**). To quantify the necessity of each DHS for *Sox2* expression, we delivered a series of LCR payloads to LP-LCR mESCs and analyzed expression of the BL6 *Sox2* allele (**Fig. 3b**). Delivery of the WT LCR completely rescued *Sox2* expression. Analysis of single and multiple DHS deletions delineated a core LCR region comprising DHSs 23-26. Within this region, deletion of DHS24 alone critically ablated *Sox2* expression to just 42% of WT, while deletion of DHS23 or DHS26 showed a lesser but significant impact. Within this core LCR, only deletion of CTCF25 was tolerated with little effect. DHS deletions outside of the core LCR showed little or no effect.

**Fig. 3.**
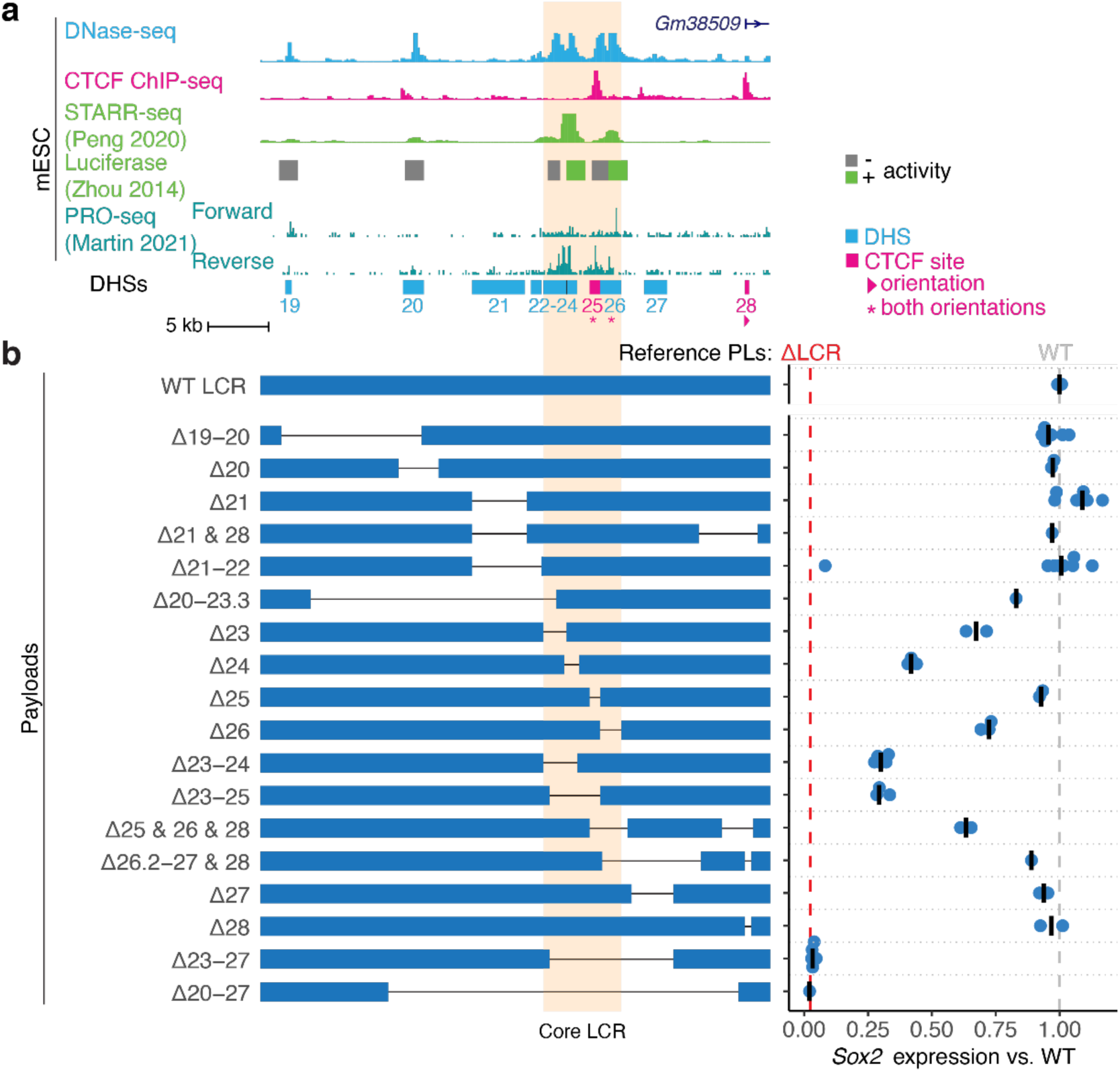
Quantifying the essentiality of *Sox2* LCR DHSs. **a.** Schematic of the *Sox2* LCR showing DNase-seq, CTCF ChIP-seq, reporter assay (STARR-seq^26^ and luciferase^17^), and PRO-seq^31^ data from mESCs. **b.** *Sox2* expression analysis for payloads with deletions of single and multiple DHSs within the core LCR delivered to LP-LCR mESCs. Blue rectangles demarcate genomic regions included in each payload. Each point represents the expression of the engineered *Sox2* allele in an independent mESC clone. Bars indicate median. Expression was scaled between 0 (*ΔSox2*) to 1 (*WT LCR*).

To investigate whether individual DHSs within the LCR function in isolation or rely on their surrounding DHSs, we delivered minimal payloads containing subsets of LCR DHSs to LP-LCR mESCs and measured their activities (**Fig. 4a** and **Supplementary Fig. S3**). We observed that a minimal payload containing the four core LCR DHSs (*23-26*) recapitulated 88% of the activity of the full WT LCR. Activities of single DHSs varied substantially: DHS24 and DHS26 were able to restore 30% and 14% of *Sox2* expression, respectively. Linking 2 copies of DHS24 (*2×24*) nearly doubled expression relative to a single copy, demonstrating that increased dosage of DHS24 without addition of any new TF activity can increase expression.

**Fig. 4.**
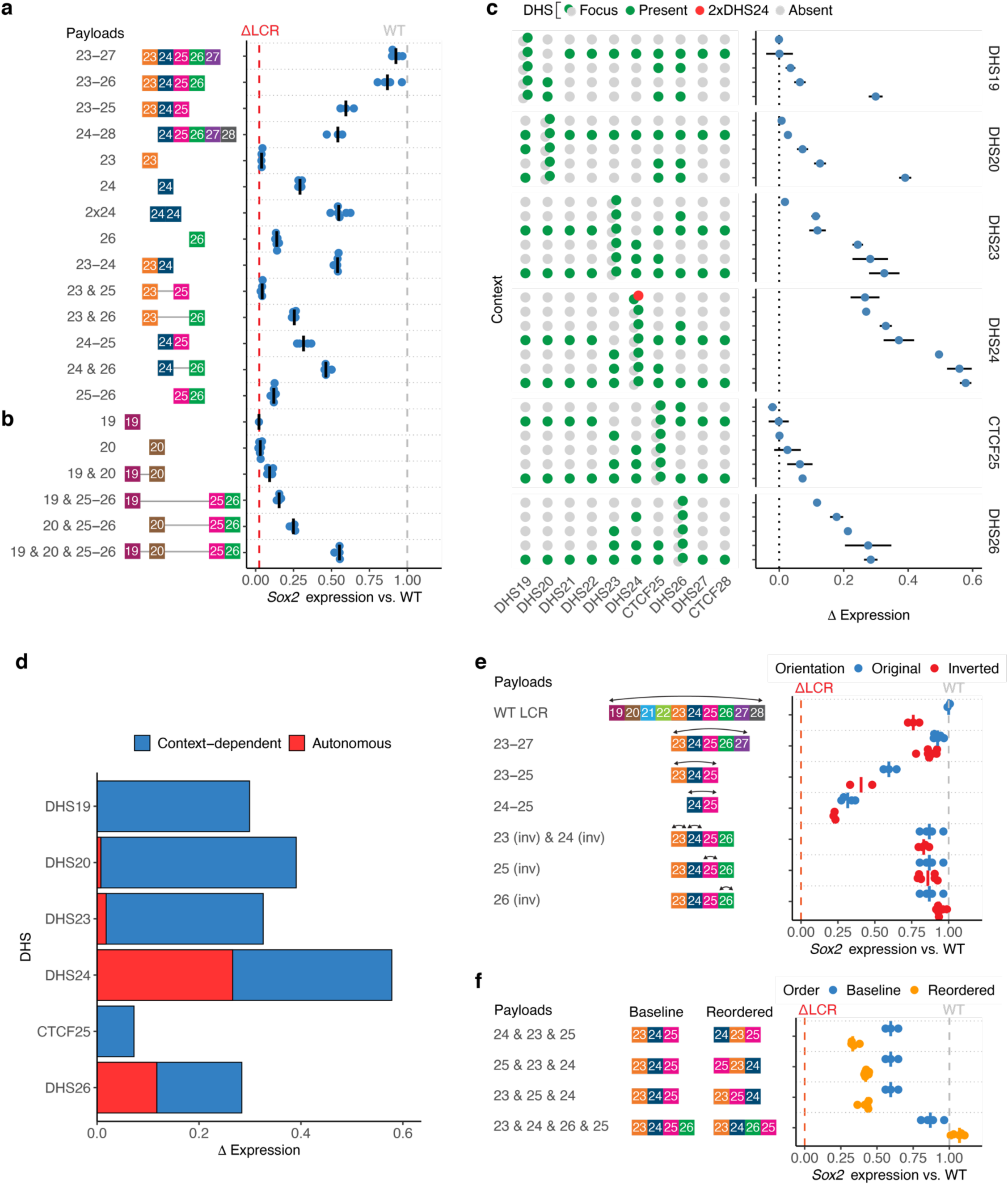
DHS function at the *Sox2* LCR is context-dependent. *Sox2* expression analysis for selected LCR DHSs replacing the full LCR; payload structures are shown in **Supplementary Fig. S3**. Each point represents the expression of the engineered *Sox2* allele in an independent mESC clone. Bars indicate median. Expression was scaled between 0 (*ΔSox2*) to 1 (*WT LCR*). **a-b**. Sufficiency of minimal payloads. **c.** Contribution of core LCR DHSs in different contexts. ΔExpression was computed as the difference in expression between payload pairs differing solely by the presence of each focus DHS. The presence (green dots) or absence (gray dots) of each DHS is indicated; two copies of DHS24 are shown in red. Points indicate mean and bars indicate SD across all pairwise combinations of clones. **d.** Summary of activity and context sensitivity of core LCR DHSs. Overall activity was defined as the maximum and context-dependent activity as the range in **c**. **e-f**. Effect of DHS orientation (**e**) and order (**f**). Expression of corresponding baseline payloads is repeated in blue for each payload. Arrows indicate extent of inversion(s) in **e**.

However, not all core LCR DHSs were sufficient to activate *Sox2* expression. DHS23, despite being required for full *Sox2* expression in the context of the entire LCR, was completely inactive on its own. But when DHS23 was linked to either DHS24 or DHS26, it augmented their overall activity nearly twofold. Deletion of DHS19 and DHS20, which were previously reported as inactive in a reporter assay^17, 26^, showed no effect on expression (**Fig. 3b**), (**Fig. 3a**). DHS19 and DHS20 also each showed no activity by themselves (**Fig. 4b**). But similar to DHS23, linking DHS20 with CTCF25 and DHS26 (*20 & 25-26*) yielded robustly increased expression (25% of WT) above that of CTCF25 and DHS26 alone (*25-26*). While DHS19 alone added little activity to CTCF25 and DHS26, it nearly doubled activity when DHS20 was present (*19 & 20 & 25-26*). Thus while DHS19, DHS20, and DHS23 can have a potent effect when linked to other enhancers, their activity is entirely context-dependent.

Given the high degree of context dependence we observed, we developed an approach to visualize the extent to which DHSs depend on their context. We identified pairs of payloads differing solely by the presence of key LCR DHSs. We calculated the contextual contribution of each of these focus DHSs as the difference in *Sox2* expression between each payload pair averaged across mESC clones (**Fig. 4c**). All DHSs exhibited marked dependence on their surrounding context. We further summarized DHS function by defining the context-dependent contribution as the portion of activity (ΔExpression) that varies between contexts, and the autonomous contribution as the remaining activity constant across all contexts (**Fig. 4d**). Partitioning DHS activity along these lines clearly highlighted the distinction between the entirely context-dependent DHS19, DHS20, and DHS23 and the more autonomous DHS24 and DHS26. Thus we conclude that the DHSs comprising the LCR are not redundant and many exhibit unexpected function when placed in novel contexts.

Context dependence might further manifest as differences in expression for different configurations of the same set of DHSs. Since certain genomic features have an intrinsic polarity, such as transcription or the interaction between cohesin and CTCF, we investigated the effect of DHS orientation on LCR activity by comparing pairs of payloads that differ in the orientation of single or multiple DHSs (**Fig. 4e**). While inversion of the entire LCR resulted in a 24% decrease in its activity, inversions of smaller LCR regions had a lesser effect. Individual inversions of DHSs 23, 24, 25, and 26 had almost no effect on expression. This is consistent with a prior report showing that surgical inversion of a CTCF recognition sequence within CTCF25 had no effect on *Sox2* expression^22^. To explore whether the effect of larger inversions might be mediated by positional differences rather than the orientation of specific TF or CTCF recognition sites, we profiled multiple permutations of the core DHS order (**Fig. 4f**). Each permutation of DHSs 23, 24, and 25 order reduced overall activity, whereas relocating CTCF25 downstream of DHS26 resulted in increased activity. These results suggest that the relative position rather than the orientation of DHSs within the LCR plays a role in their function.

Finally, to investigate whether individual DHSs may be functionally specialized, we examined published ChIP-seq and ChIP-nexus data over the LCR^4, 27–30^ (**Supplementary Fig. S3**). We noted highly similar occupancy patterns across all DHSs: most DHSs were occupied by Nanog, Oct4, Klf4, Esrrb, and Zic3, with Sox2 and Pbx binding to a more restricted set of sites. Only DHS23 and DHS24 exhibited robust activity in reporter assays. DHS24 uniquely showed a high level of transcription initiation using PRO-seq^31^, oriented towards the *Sox2* gene. These data show that individual DHSs each manifest characteristic regulatory signatures, but no single feature explains the distinction between the context-dependent and autonomous DHSs.

### Modeling the regulatory architecture of the *Sox2* locus

To coherently summarize the architecture of the *Sox2* locus, we investigated linear regression models that predict *Sox2* expression from surrounding DHS composition and configuration. We established a baseline model considering only key proximal regions along with the four DHSs in the core LCR. Performance improved substantially upon incorporation of interactions among core LCR DHSs, in particular the contribution of DHS23 in conjunction with DHS24 or DHS26, and CTCF25 in the presence of DHS26 (**Fig. 5a**). Performance was higher when restricting to these three top interaction indicators than when considering all pairwise interactions. Given the potential influence of DHS configuration on function, we extended our top interaction regression model to include the orientation of each core LCR DHS and relative order encoded by indicators for DHS24 preceding DHS23 or CTCF25 preceding DHS24. Inclusion of either orientation or order separately improved model performance, but a model including order alone provided a better fit than orientation alone or a combination of both (**Fig. 5a**). The coefficients of the best fit model showed strong weights for the presence of proximal regulatory regions and core LCR DHSs, as well as key interactions and DHS order (**Fig. 5b**). While most payloads were predicted with little error, we noted that the contribution of DHS26 as well as inversions and permuted constructs demonstrated consistently higher error than other constructs, suggesting additional context effects not fully captured by our simple model (**Supplementary Fig. S4**). This model demonstrates the power of a synthetic regulatory genomics approach to dissect locus architecture in the presence of substantial context dependence.

**Fig. 5.**
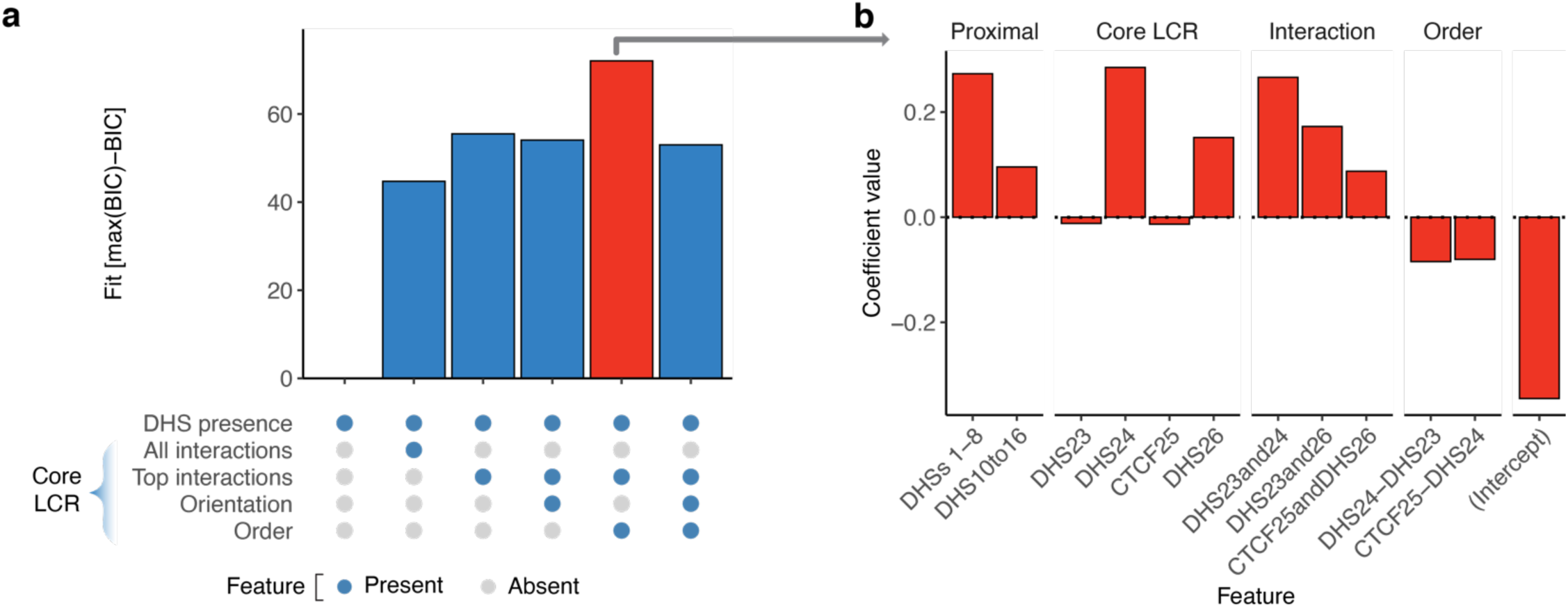
Modeling the regulatory architecture of the *Sox2* locus. **a.** Relative predictive performance of linear regression models for *Sox2* expression. Models included features for DHS presence throughout the locus and orientation, order, and interaction terms for core LCR DHSs as indicated by blue dots. Model performance was measured by Bayesian Information Criteria (BIC) and presented as Fit (difference from maximum BIC). Red indicates the model with best fit. **b.** Coefficients for the best model.

### TF-scale dissection of the core *Sox2* LCR

Given the complex interdependencies among core LCR DHSs when considered as units, we used synthetic DNA to investigate function at the level of individual TF recognition sequences. We designed base payloads covering the core LCR DHSs that avoided or shortened a handful of repetitive regions to facilitate chemical synthesis (**Fig. 6a**). We then designed a series of derivative payloads including surgical deletions, mutations or inversions to assess the individual and collective functions of putative TF or CTCF binding sites identified through analysis of TF motif matches and *in vivo* DNA accessibility and TF occupancy data (**Fig. 6b**).

**Fig. 6.**
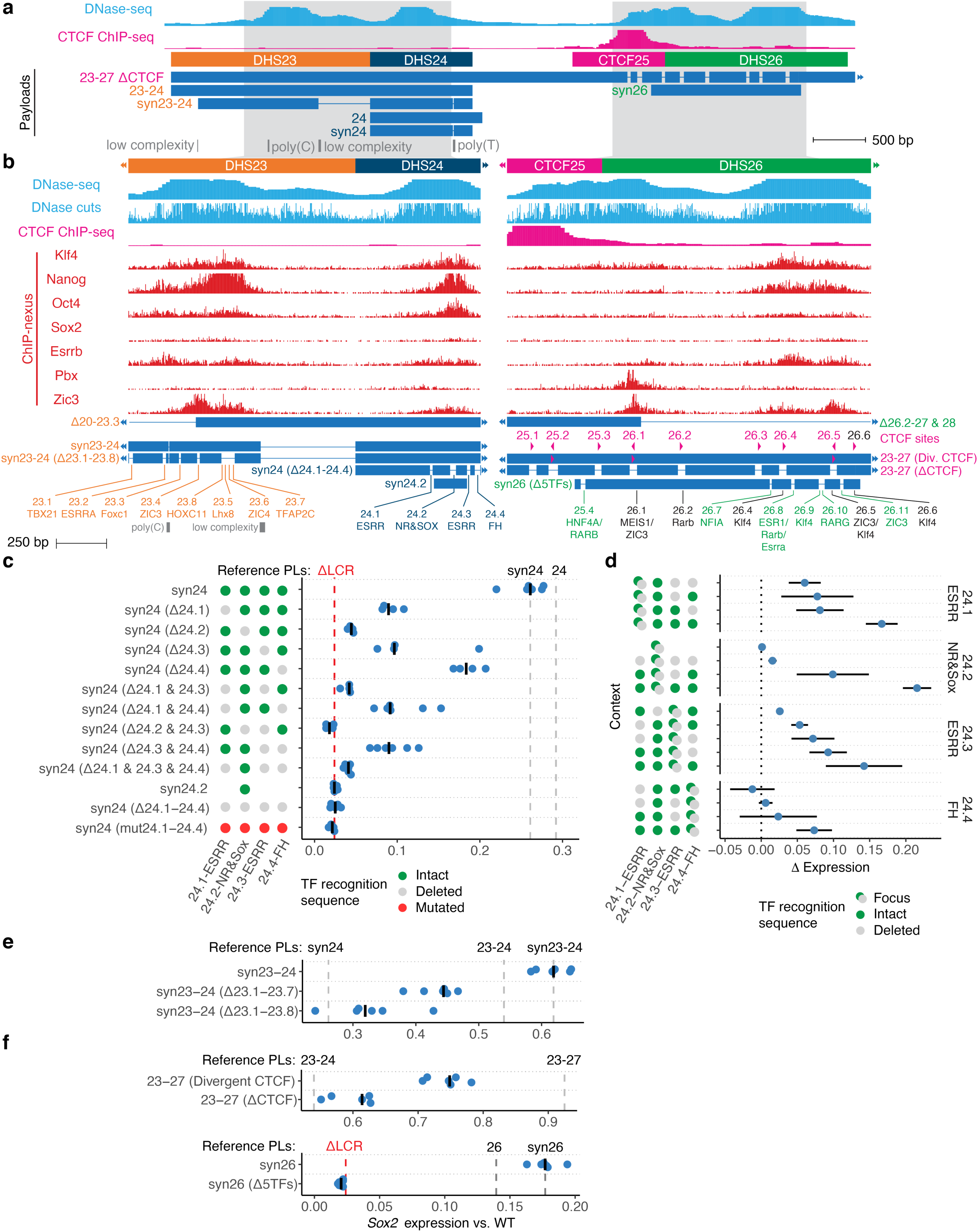
TF-scale dissection of LCR function. **a.** Genomic region surrounding the *Sox2* LCR DHSs 23-26 showing DNase-seq and CTCF ChIP-seq data in mESCs and selected payloads. Gray rectangles indicate poly(C) and poly(T) sequences shortened in synthetic (syn) versions of DHSs 23 or 24. **b.** Enlargement of engineered regions within DHS23 to DHS24 (left) and CTCF25 to DHS26 (right) showing DNase-seq windowed density and per-nucleotide cleavages, and ChIP-nexus data for selected TFs in mESCs. Payload schemes indicate surgical deletions of predicted TF recognition sites. Magenta triangles denote CTCF sites and their native orientation. Additional TF sites overlapping CTCF sites are shown in black. **c-f**. *Sox2* expression analysis for perturbations of TF recognition sequences within the core LCR. Payloads include deletions (Δ) of TF or CTCF recognition sequences shown in **b**. Each point represents the expression of the engineered *Sox2* allele in an independent mESC clone relative to baseline payloads (dashed vertical lines). Bars indicate median. Expression was scaled between 0 (*ΔSox2*) to 1 (*WT LCR*), and vertical gray lines indicate median expression of relevant baseline payloads. **c.** Analysis of DHS24. *syn24.2* contains only a minimal region surrounding DHS24.2 (see **b**). *syn24 (mut24.1-24.4)* contains point substitutions that ablate TF recognition sequences 24.1-24.4 instead of deletions (**Supplementary Fig. S5**). **d.** Context-sensitive activity of TF sites 24.1-24.4. ΔExpression was computed as the difference in expression between payload pairs differing solely by the presence of each focus TF recognition sequence. The presence (blue dots) or absence (gray dots) of each TF site is indicated. Points indicate mean and bars indicate SD across all pairwise combinations of clones. **e.** Analysis of DHS23. **f.** Analysis of CTCF25 and DHS26. *23-27 (ΔCTCF)* has 8 CTCF sites surgically deleted. *23-27 (Divergent CTCF)* has surgical inversion of 3 CTCF sites (TF sites 25.2, 26.1 and 26.5) so that all 9 CTCF sites lie in divergent orientation relative to *Sox2*.

We started with DHS24, which is the strongest single DHS in the LCR and is itself capable of reproducing 30% of WT LCR activity (**Fig. 4a**). We identified four distinct TF recognition sequence sites, numbered 24.1-24.4 (**Fig. 6b** and **Supplementary Fig. S5**). These sites were predicted to be occupied by key mESC regulators, including the ESRR, nuclear receptor, SOX, and forkhead TF families. Deletion or mutation of all four sites was sufficient to fully ablate activation of *Sox2* by *syn24* (**Fig. 6c**). All single deletions showed at least some effect, ranging from an 83% (24.2) to 28% (24.4) reduction of base activity (**Fig. 6c**). Double or triple deletions showed a further negative effect, essentially reaching null when 24.2 and 24.3 were both deleted. A minimal payload containing the most essential site based on the deletion analysis (24.2) with its flanking regions was unable to confer any *Sox2* expression. We analyzed the contribution to *Sox2* expression of each TF site relative to its context, comparing pairs of payloads differing by the presence of a focus TF site (**Fig. 6d**). This analysis identified a clear context dependence for all four TF sites, and the contribution of each TF site increased with the number of other sites present.

As DHS23 alone is incapable of activating *Sox2* expression (**Fig. 4a**), it was analyzed linked to DHS24 (*syn23-24*, **Fig. 6b**). The activity of this payload was slightly higher than the comparable non-synthetic payload including DHSs 23-24 (**Fig. 6e**), possibly due to the decreased distance between the core DHS peaks of DHS23 and DHS24 in the synthetic configuration. Deletion of 8 recognition sequence sites within DHS23 reduced activity by 47%, approaching that of *syn24* alone and suggesting that they are the key sequences responsible for DHS23 function.

This synthetic approach further permitted assessment of two conserved DHSs from the human *SOX2* locus, orthologous to mouse DHS23 and DHS24 (**Supplementary Fig. S6a**). While mouse DHS23 and DHS24 combined are sufficient to confer over half of WT *Sox2* expression, the two orthologous human DHSs surprisingly showed no activity whatsoever (**Supplementary Fig. S6b**). While the core of the human DHSs showed sequence conservation with mouse sequence, analysis of TF recognition sequences showed a marked divergence relative to the orthologous mouse sites. This suggests that, despite the sequence orthology, the human LCR has diverged mechanistically.

Finally, we investigated CTCF25 and DHS26, which delineate a region covered by continuous DNA accessibility that comprises two broad DHS peaks positioned closely together (**Fig. 6a-b**). As redundant CTCF binding is a feature of other insulator elements^19, 32, 33^, we performed a search for motif matches in addition to the previously reported CTCF site 25.2^22^. This identified 9 CTCF recognition sequences in CTCF25 and DHS26, 3 of which are in convergent orientation relative to *Sox2* (**Fig. 6b** and **Supplementary Fig. S5**). While many of these sites lie outside the primary ChIP-seq peak, they reside within a DHS and demonstrate a trace level of CTCF occupancy in WT cells (**Fig. 6b**), so we reasoned that they might be partially redundant with 25.2. To test this hypothesis, we investigated whether inversion or deletion of multiple sites might result in a stronger phenotype than the reported inversion of 25.2 alone. Inversion of the 41-bp CTCF footprint at the 3 sites in convergent orientation to *Sox2* in a baseline payload incorporating DHS23 through DHS27 (*23-27 (Divergent CTCF)*, **Fig. 6b**) led to a 20% reduction in expression (**Fig. 6f**). Deletion of 8 CTCF recognition sequences (*23-27 (ΔCTCF)*, **Fig. 6b**) led to an even greater reduction of 36% (**Fig. 6f**), bringing activity down nearly to the baseline of the remaining DHS23 and DHS24, and showing that these sites are collectively required for full function of the CTCF25-DHS26 segment.

As DHS26 itself can function as an enhancer (**Fig. 4a**), we next investigated the function of TF sites not overlapping with CTCF recognition sequences. A synthetic DHS26 payload demonstrated slightly higher activity than the non-synthetic DHS26 (**Fig. 6f**), likely because it extended into CTCF25 to include a predicted HNF4A/RARB site (25.4) and a CTCF site (25.3) (**Fig. 6b** and **Supplementary Fig. S5**). Deletion of 5 TF recognition sequence sites in this context also completely abrogated activity (**Fig. 6d**). Finally, *Δ26.2-27 & 28*, while lacking all 5 of these TF sites and CTCF sites 26.2-26.6, but containing 26.1 (a CTCF site also occupied by Pbx, and Zic3), showed nearly full expression (**Fig. 6b** and **Supplementary Fig. S6c**). This suggests that, like DHS24, CTCF25-DHS26 is a complex regulatory element harboring multiple essential TF sites whose binding is context dependent and synergistic.

## Discussion

Dissection of the *Sox2* LCR shows it to be a highly complex and species-specific element whose function depends on the specific conformation of its constituent DHSs. Sufficiency analysis shows that the contribution of an individual DHS depends on its surrounding context, and DHS activity ranges from complete autonomy to completely context-dependent. The identification of three fully context-dependent enhancers at the same locus, DHS19, DHS20, and DHS23, enables comparison among them: Unlike DHS23 which was essential for full activity of the LCR, DHS19 and DHS20 were dispensable at their native locations, where it is possible that distance moderates their influence on DHSs in the core LCR. DHS19 differed from DHS20 and DHS23 in that it required DHS20 to robustly augment DHS26, suggesting an additional level of context dependence. The combination of the two context-dependent enhancers DHS19 and DHS20 produced activity only just above zero, suggesting that robust function requires the presence at least one autonomous enhancer. DHS23 augmented the activity of both DHS24 and DHS26, suggesting its function is not strictly tied to a single partner and at least somewhat flexible. The behavior of these DHSs resembles those recently identified at the *Fgf5* and *Hba* loci^38, 39^, suggesting that context-dependent DHSs are widespread throughout the genome. Finally, these context dependencies were not recapitulated in reporter assays or more limited engineering of the endogenous locus, underscoring the importance of a synthetic regulatory genomics approach for comprehensive analysis of regulatory architecture in context.

The *Sox2* LCR demonstrates a high degree of cooperativity both at the level of individual TF sites as well as DHSs, but our results do not address whether this arises from direct TF-TF interactions, indirect interaction mediated through the chromatin template, local TF concentrations, or other mechanisms^40, 41^. Deletions of TF recognition sites had larger effects in more complete contexts, which contrasts with prior results showing that the effect of point variants is buffered by stronger DHSs^42^. We speculate that this discrepancy is related to the size of the perturbation, as TF binding can more readily tolerate point changes without falling below the threshold needed for activity^43^. As Sox2 binds indirectly at many of the LCR DHSs (**Supplementary Fig. S3**), it is also possible that this context effect is partially mediated through altered Sox2 levels in trans. We expect that the synthetic payloads containing surgical TF deletions (**Fig. 6**) will provide a roadmap for investigation of the sequence determinants of these context effects and their effect on the regulatory complexes involved.

Our approach investigates function at the endogenous locus, preserving its long-distance architecture and surrounding genomic context. The WT constructs replacing the full locus or just the LCR were sufficient to rescue expression in mESC without any detectable defect, but it is possible their chromatin state might still differ from the endogenous locus. The payloads themselves are epigenetically naïve in that they do not pass through development and their chromatin state is established after delivery based on the surrounding context. Characteristic chromatin and expression patterns are readily reestablished on DNA that passes through the germline^34, 35^, but it is possible that DNA transfected to cycling cells exhibits only partial functionality^36, 37^. For example, the failure of DHS23 to activate *Sox2* expression alone might arise from an inability to independently establish a functional accessible state. We expect future studies might study the establishment of chromatin state on rewritten genomic loci in more detail.

LCR DHSs might act together differently in contexts beyond mESCs in culture. For example, while DHS19 and DHS20 lie outside the core LCR and are dispensable in mESCs, their potent effect when linked more closely to the core LCR suggests a possible latent function relevant to other cell states. Similarly, while *Sox2* expression for a given payload was highly consistent across replicate clones, a single clone lacking DHSs 21-22 showed a nearly complete absence of activity in contrast to the other 8 clones which showed near-WT activity (**Fig. 3b**). This outlier clone passed extensive genotyping and Capture-seq to demonstrate that the locus is intact, suggesting that seemingly redundant sites may influence robustness in certain circumstances.

We have described a general approach to dissect locus architecture in-place from 100-kb scale down to individual bp-level TF binding events, and have shown that analyses of sufficiency and combinatorial rearrangements at the endogenous locus reveal unexpected behavior relative to reporter assays or single deletion analyses. We have already employed Big-IN in multiple cell types and loci^25^, suggesting that our approach is readily generalizable. While large-scale multiplexed CRISPR screens typically do not verify the effect of each edit, our genotyping and genomics validation has shown that a recombinase-based approach avoiding double stranded breaks yields high efficiency delivery with little or no off-target integration. We expect that engineering of large or complex payloads in yeast will become more widely available. In parallel, Golden Gate-based cloning of shorter payloads from commercial synthetic DNA provides a rapid, scalable, and accessible strategy for high-throughput analyses. These developments suggest that synthetic regulatory genomics approaches will rapidly be deployed to investigate function across additional contexts and loci.

## Methods

### Cloning landing pad and CRISPR/Cas9 plasmids for genome integration

pLP-PIGA (pLP140) (Addgene #168461) was described previously^25^ and harbors a pEF1α-mScarlet-P2A-CreERT2-P2A-PuroR-P2A-hmPIGA-EIF1pA cassette flanked by loxM and loxP sites.

pLP-PIGA2 (pLP300) (Addgene #168462) (**Fig. 1a**) was described previously^25^ and harbors a pEF1α-PuroR-P2A-hmPIGA-P2A-mScarlet-EIF1pA cassette flanked by loxM and loxP sites, as well as a pPGK1-ΔTK-SV40pA backbone counter-selectable marker cassette.

To derive LP-PIGA3 (pLP305) (**Fig. 1b**), an intermediate pLP303a was cloned by removing mScarlet-P2A-CreERT2-P2A from pLP-PIGA using NcoI and SalI digest followed by a fill-in reaction with Klenow DNA Polymerase and self-ligation to yield a minimal LP plasmid consisting of pEF1α-PuroR-P2A-hmPIGA-EIF1pA cassette flanked by loxM and loxP sites (**Supplementary Table S5**). Sleeping Beauty inverted terminal repeats (ITRs) were amplified from pMH005^44^ using primers oRB_277 + oRB_278 for ITR(L) and oRB_279 + oRB_287 for ITR(R). LP-PIGA3 was cloned using a BsaI Golden Gate reaction in which ITRs were cloned outside the LP region of pLP303a.

To target LPs to specific genomic loci, guide RNAs (gRNAs) (**Supplementary Table S8)** were cloned into pSpCas9(BB)-2A-Puro (pCas9-Puro, Addgene #62988) or pSpCas9(BB)-2A-GFP (pCas9-GFP, Addgene #48138) plasmids using BbsI Golden Gate reactions as described^45^. LP homology arms (HAs) corresponding to the genomic sequence flanking the Cas9 cut sites were amplified from a BAC (**Supplementary Table S6)**. HAs were cloned flanking the LoxM and LoxP sites in pLP-PIGA2 using a BsaI Golden Gate reaction or flanking the ITRs in pLP-PIGA3 using a BsmBI Golden Gate reaction (**Supplementary Table S7**). DNA suitable for transfection was prepped using the ZymoPURE maxiprep kit (Zymo Research D4203) according to the manufacturer’s protocol.

Landing pad sequences (excluding HAs and backbone) are provided in **Data S1**.

### Payload assembly

All payloads were assembled into pLM1110^25^ (Addgene #168460) or pRB051 (a derivative of pLM1110). Both are multifunctional YAC/BAC (yeast and bacterial artificial chromosomes) vectors supporting low-copy DNA propagation, selection (LEU2), and efficient homology-dependent recombination in yeast; low-copy propagation, selection (KanR), and copy number induction in TransforMax EPI300 *E. coli*^46^, and transient visualization and selection (eGFP-BSD, enhanced green fluorescent protein - Blasticidin-S deaminase) in mammalian cells^25^.

Four different strategies were used to assemble payloads into these YAC/BACs (**Supplementary Table S1, Data S1**), using different primers (**Supplementary Table S5)**, gRNAs (**Supplementary Table S8),** and synthetic DNA fragments (**Supplementary Table S9**):

#### 1) Yeast CRISPR/Cas9 Assembly

Desired segments of *Sox2* BAC RP23-144O8 were released by an *in vitro* CRISPR/Cas9 digestion using a pair of synthetic gRNAs and recombinant Cas9 and assembled into BsaI-digested pLM1110 as previously described^25^.

#### 2) Yeast Assembly

PCR amplicons or synthetic DNA (IDT) tiling the desired payload with >75 bp overlap were assembled into pLM1110. Yeast cells were transformed with 20-50 ng I-SceI-digested pLM1110, 100 ng of each DNA fragment, and 50 ng terminal linker fragments (∼400 bp gBlocks, IDT) to enable homologous recombination-dependent assembly followed by selection for LEU^+^ phenotype.

#### 3) Yeast CRISPR/Cas9 Editing

Existing payloads were modified in yeast using CRISPR/Cas9 and synthetic linker fragment(s) (IDT) to mediate up to two deletions, insertions, or inversions simultaneously^47^:

i. A yeast strain carrying a parental payload YAC/BAC was pre-transformed with 100 µg SpCas9 expression vector pNA0519 (carrying a ScHIS3 marker) or pCTC019 (carrying a SpHIS5 marker), and selected for a LEU^+^/HIS^+^ phenotype. Yeast were then transformed again with 100 µg single/dual gRNA expression plasmid pNA0304/pNA0308^48^ carrying a URA3 marker and with 100 µg synthetic linker fragment(s), and selected for a LEU^+^/HIS^+^/URA^+^ phenotype. gRNAs (**Supplementary Table S8)** were cloned into the single/dual gRNA yeast expression plasmids pNA304/pNA0308, respectively, using *Hind*III and/or *Not*I Gibson Assembly reactions as described^48^. Yeast Cas9 expression vector pNA0519 pRS413-TEF1p-Cas9 was derived from pNA0306^49^ by subcloning the TEF1p-Cas9-CYC1t cassette into a pRS413 (ATCC 87518) in order to replace the LEU2 marker with HIS3^50^.
ii. Alternatively, yeast cells were transformed with pYTK-Cas9 plasmids (carrying HIS3 marker) that co-express SpCas9 and a gRNA and with synthetic linker fragment(s), and selected for LEU^+^/HIS^+^ phenotype. gRNAs (**Supplementary Table S8)** were cloned into pYTK-Cas9^47^ using a BsmBI Golden Gate reaction.

#### 4) Golden Gate Assembly (GGA)

DNA fragments were assembled into pLM1110 using BsaI or into pRB051 using Esp3I. pRB051 was cloned by PCR-amplifying the RFP transcriptional unit from pLM1110 using primers oRB_564 + oRB_565, digesting the product with Esp3I, and performing a BsaI Golden Gate Assembly into pLM1110. For payload assembly, 100 ng of pLM1110 or pRB051 was mixed with 20 ng of each DNA fragment containing terminal BsaI or Esp3I sites, respectively, designed to mediate assembly with neighboring fragments (DNA fragments were sourced either from synthetic fragments (IDT) or from existing payloads by PCR-amplification), and with 0.4 µL BsaI-HF V2 (NEB R3733S) or Esp3I (NEB R0734S), 1.5 µL 1 mg/mL BSA, 1 µL T4 Quick Ligase (NEB M2200S) and 1.5 µL T4 Ligase Buffer (NEB) in a total volume of 15 µL. Reactions were cycled 25 times between 37°C (3 min) and 16 °C (4 min), heat inactivated at 50 °C (5 min) and 80 °C (5 min) before transforming into TransforMax EPI300 cells.

### Yeast transformation, payload validation and transfer to *E. coli*

Yeast transformations were performed using the Lithium acetate method^51^ with *Saccharomyces cerevisiae* strain BY4741. For screening of correct clones, DNA was isolated from individual yeast colonies by resuspension in 10-40 µl of 20 mM NaOH and boiling for 3 cycles of 95 °C for 3 min and 4°C for 1 min. 2 µl of yeast lysate was used as a template in a 10 µl GoTaq Green reaction (Promega M7123) with 0.25 µM of primers. Clones were screened for the intended payload structure using primers that target newly formed junctions (positive screen), and in some cases using primers that target regions that are present in the parental plasmid but absent in the intended plasmid (negative screen). YAC/BACs were isolated from candidate yeast clones using the Zymo yeast miniprep I protocol (Zymo Research D2001) and transformed into TransforMax EPI300 *E. Coli* cells by electroporation using the manufacturer’s protocol (Lucigen EC300150).

### Payload DNA screening and preparation

For initial verification, TransforMax EPI300 *E. coli* colonies were picked into 3-5 mL LB-Kan supplemented with CopyControl Induction Solution (Lucigen CCIS125) and cultured overnight at 30°C with shaking at 220 RPM. YAC/BACs smaller than 20 kb were isolated using the Zyppy Plasmid Miniprep Kit (Zymo Research D4020).

For larger YAC/BACs, crude DNA extraction was performed as follows: 2-3 mL of induced culture was spun down, resuspended with 300 µL RNase A-supplemented Buffer P1 (QIAGEN 19051), and topped with 300 µL of Buffer P2 (QIAGEN 19052). The suspensions were inverted 10 times, incubated at room temperature for 2 min, topped with 300 µL Buffer P3 (QIAGEN 19053), inverted 10 times and spun down at 12000 RPM for 5 min. Supernatants were transferred to new 2 mL tubes, topped with 900 µL isopropanol, mixed by inverting, and spun down at 12000 RPM for 5 min. Supernatants were discarded, pellets were washed with 500 µl of 70% EtOH, and spun down at 12,000 RPM for 1 min. Supernatants were discarded and the DNA pellets were air-dried and resuspended in 30 µL TE buffer. Tubes were spun down at 12000 RPM for 1 min and clear supernatants transferred to new 1.5 mL tubes.

Payload assembly verification was performed by diagnostic digestion with restriction enzymes, PCR verification of new junctions, and Sanger sequencing, depending on the payload. All YAC/BACs were subsequently sequenced to high coverage depth.

Sequence-verified YAC/BAC clones were grown overnight in 2.5 mL cultures of LB-Kan at 30 °C with shaking, diluted 1:100 in LB-Kan supplemented with CopyControl Induction Solution, and incubated for an additional 8-16 hours at 30 °C with shaking. For transfection, DNA was purified from induced *E. Coli* using the ZymoPURE maxiprep kit (Zymo Research D4203) for YAC/BACs <20 kb or using the Nucleobond XtraBAC kit (Takara 740436) for YAC/BACs >20 kb. YAC/BAC preps were stored at 4 °C.

### mESC culture

C57BL6/6J × CAST/EiJ (BL6xCAST) mESCs were cultured as described^25^. Specifically, mESCs were cultured on plates coated with 0.1% gelatin (EMD Millipore ES006-B) in 80/20 medium comprising 80% 2i medium and 20% mESC medium. 2i medium contained a 1:1 mixture of Advanced DMEM/F12 (ThermoFisher 12634010) and Neurobasal-A (ThermoFisher 10888022) supplemented with 1% N2 Supplement (ThermoFisher 17502048), 2% B27 Supplement (ThermoFisher 17504044), 1% GlutaMAX (ThermoFisher 35050061), 1% Pen-Strep (ThermoFisher 15140122), 0.1 mM 2-mercaptoethanol (Sigma M3148), 1,250 U/mL LIF (ESGRO ESG1107l), 3 µM CHIR99021 (R&D Systems 4423), and 1 µM PD0325901 (Sigma PZ0162). mESC medium contained KnockOut DMEM (ThermoFisher 10829018) supplemented with 15% FBS (BenchMark 100-106), 0.1 mM 2-mercaptoethanol, 1% GlutaMAX, 1% MEM nonessential amino acids (ThermoFisher 11140050), 1% nucleosides (EMD Millipore ES008-D), 1% Pen-Strep, and 1,250 U/mL LIF. Cells were grown at 37 °C in a humidified atmosphere of 5% CO2 and passaged on average twice per week. MK6 C57BL/6J mESCs were provided by the NYU Rodent Genetic Engineering Laboratory.

### mESC genome engineering

Landing pad integrations were performed (**Supplementary Table S3)** using the Neon Transfection System as previously described into BL6xCAST Δ*Piga* mESCs, in which the endogenous *Piga* gene was deleted^25^. LP-PIGA A1 mESCs, in which LP-PIGA replaces the BL6 *Sox2* allele were previously described^25^.

LP-PIGA2 integration at *Sox2* (replacing a 143-kb genomic region) was performed using 5 µg of pLP-PIGA2 and 2.5 µg of each pCas9-GFP plasmid expressing BL6-specific gRNAs mSox2-5p-1 and mSox2-3p-5, followed by 1 µg/mL puromycin (ThermoFisher A1113803) selection for LP-harboring mESCs and 1 µM ganciclovir (GCV, Sigma PHR1593) selection against LP-PIGA2 backbone’s HSV1-ΔTK gene.

LP-PIGA3 integration at the *Sox2* LCR (replacing a 41-kb genomic region) was performed similarly with pCas9-Puro plasmids expressing the non-allele specific gRNA mSox2-DHS18-19 and the BL6-specific gRNA mSox2-3p-5, followed by 1 µg/mL puromycin selection. PCR genotyping and Capture-seq validation were used to identify clone LP-LCR H1 (**Supplementary Fig. S1b** and **Supplementary Fig. S1d**), to which a few LCR payloads were delivered (**Supplementary Table S3**). DELLY analysis later identified a 7.7 kb deletion (**Supplementary Fig. S1d**) in the CAST allele surrounding the mSox2-DHS18-19 gRNA site in a small number of payload clones derived from LP-LCR H1 cells (not included in this study), indicating a mixed population of cells in the original LP-LCR culture. We therefore sub-cloned LP-LCR H1 to isolate clone LP-LCR C1 and confirmed it did not harbor the CAST allele deletion (**Supplementary Fig. S1 c-d**). LP-LCR C1 was used for all subsequent LCR deliveries.

Payload deliveries were performed as previously described^25^ with 1-5 × 10^6^ mESCs, 1-10 µg YAC/BAC payload DNA and 2-5 µg pCAG-iCre plasmid (Addgene #89573) per transfection, depending on the payload size (larger payloads required more cells, PL DNA and pCAG-iCre plasmid to obtain sufficient correct mESC clones). Transfected mESCs were selected with 10 µg/mL blasticidin for 2 days starting day 1 post-transfection and with 2 nM proaerolysin for 2 days starting day 6 or 7 post-transfection.

For both LP integration and payload delivery, approximately 9 days post-transfection, individual mESC clones were manually picked into gelatinized 96-well plates prefilled with 100 µL 80/20 media. Two days post-picking, clones were replicated into two gelatinized 96-well plates at 90% and 10% relative densities. Three days later, crude gDNA was extracted from the 90% plate as described^25^ and used in PCR genotyping to identify candidate clones, which were then expanded from the 10% density plate for further verification and phenotypic characterization. Genomic DNA was extracted from expanded clones using the DNeasy Blood & Tissue kit (QIAGEN 69506).

### mESC genotyping

Genotyping mESC clones was performed either using PCR followed by gel electrophoresis as described^25^ or using real-time quantitative PCR (qPCR), which was performed with the KAPA SYBR FAST (Kapa Biosystems KK4610) on a LightCycler 480 Real-Time PCR System (Roche) using either 96-well or 384-well qPCR plates. In most cases loading was performed using an Echo 550 liquid handler (Labcyte): A 384-well qPCR plate was prefilled with 5 µL KAPA SYBR FAST and 4 µL water per well. 100 nL of each 100 µM primer and 0.5 µL of each crude gDNA sample were transferred by the Echo. Genotyping payload clones typically included, in addition to assays designed to detect the newly-formed left and right junctions, assays to detect the loss of the landing pad, the absence of the payload YAC/BAC backbone, and for large (>100 kb) payloads, allele-specific assays to detect delivered regions of the *Sox2* locus. Genotyping primers are listed in **Supplementary Table S10 and Supplementary Table S11**.

### High-throughput sequencing verification

Illumina sequencing libraries were prepared from purified DNA using three methods, as listed in **Supplementary Table S12:**

1. The Illumina dsDNA protocol was previously described in ^25^. 1 µg of DNA was sheared in a 96-well microplate using the Covaris LE220 (450 W, 10% Duty Factor, 200 cycles per burst, and 90-s treatment time) to yield fragments between 500 to 900 bp. DNA fragments were end-repaired with T4 DNA polymerase, Klenow DNA polymerase, and T4 polynucleotide kinase (New England Biolabs), and A-tailed using Klenow (3′-5′ exo-; New England Biolabs). Illumina sequencing adapters were then ligated to DNA ends using Quick Ligase (New England Biolabs). The post-ligation product was purified using 18% Sera-Mag Magnetic Beads (Cytiva) in polyethylene glycol. DNA libraries were amplified with KAPA 2× Hi-Fi Hotstart Readymix (Roche) and purified with 18% Sera-Mag Magnetic Beads in polyethylene glycol.
2. One-Pot dsDNA was performed as in the Illumina dsDNA protocol, except that 250 ng of sheared DNA was end-repaired and A-tailed in a single reaction using dATP, dNTPs mix, T4 DNA Polymerase, T4 Polynucleotide Kinase, and Taq Polymerase (New England Biolabs) and incubated in a thermocycler at 12 °C for 10 min, 37 °C for 10 min, and 72 °C for 20 min.
3. The NEBNext Ultra FS II kit (NEB E7805L) was used according to the provided protocol.

Final library concentrations were measured on a Qubit using the dsDNA High Sensitivity Assay Kit (Invitrogen).

Hybridization capture (Capture-seq) for targeted resequencing of engineered regions was performed as previously described^25^. Biotinylated bait was generated using nick translation from BACs covering the *Sox2* locus (RP23-144O8 or RP23-274P9; see **Supplementary Table S6**), landing pad plasmid (LP-PIGA2 or LP-PIGA3), payload plasmid backbone, pCAG-iCre and pSpCas9 plasmids. Bait sets and sequencing statistics are listed in **Supplementary Table S12**. Sequencing libraries were sequenced in paired-end mode on an Illumina NextSeq 500 operated at the Institute for Systems Genetics. Reads were demultiplexed with Illumina bcl2fastq v2.20 requiring perfect match to the indexing BC sequence. All whole-genome sequencing and Capture-seq data were processed using a uniform mapping pipeline. Illumina sequencing adapters were trimmed with Trimmomatic v0.39^52^. Reads were aligned using BWA v0.7.17^53^ to the appropriate reference genome (GRCm38/mm10 or GRCh38/hg38), including unscaffolded contigs and alternate references, as well as to independent custom references for relevant vectors. PCR duplicates were marked by samblaster v0.1.24^54^. Per-base coverage depth tracks were generated using BEDOPS v2.4.4^55^.

Variant calling was performed using a standard pipeline based on bcftools v1.14 ^56^:

bcftools mpileup --excl-flags UNMAP,SECONDARY,DUP --redo-BAQ --adjust-MQ 50 –

gap-frac 0.05 --max-depth 10000 --max-idepth 200000 -a DP,AD --output-type u |

bcftools call --keep-alts --ploidy [1|2] --multiallelic-caller -f GQ --output-type u |

bcftools norm --check-ref w --output-type u |

bcftools filter -i “INFO/DP>=10 & QUAL>=10 & GQ>=99 & FORMAT/DP>=10” --set-GTs

. --output-type u |

bcftools view -i ‘GT=“alt”’ --trim-alt-alleles --output-type z

Bcftools call --ploidy was set to 1 for custom references and 2 for autosomes in mm10.

Large deletions and structural variants were called using DELLY v0.8.7 excluding telomeric and centromeric regions^57^. Variants were required to PASS filters, to have at least 10 paired-end reads and 20% of paired-end reads supporting the variant allele. DELLY results for YAC/BAC and capture sequencing data were inspected manually and used to choose clones for successive analysis.

Data were visualized and explored using the University of California, Santa Cruz Genome Browser^58^. The full processing pipeline is available at https://github.com/mauranolab/mapping.

### Payload YAC/BAC sequence verification

To identify potential sample swaps or contamination, mean sequencing coverage was calculated for the genomic regions corresponding to the payload sequence and to the non-payload sequence, defined as the engineered *Sox2* regions that do not overlap the payload. To minimize coverage abnormalities associated with sequence termini, 400 bp were clipped from each end of continuous genomic regions. Mean coverage was normalized by the payload YAC/BAC backbone coverage. We expect a normalized coverage of 1±0.4 for the payload region and <0.1 for the non-payload region. Human orthologous sequences and regions smaller than 200 bp after clipping were ignored. Coverage analysis results and quality control (QC) calls for each sample are reported in **Supplementary Table S13** and summarized in **Supplementary Table S2**.

To detect payload DNA variants that might have been introduced during assembly or propagation in yeast or bacteria, we analyzed high confidence variant calls relative to the payload custom reference having sequencing depth above 50 and quality score above 100 (DP ≥ 50 & QUAL ≥ 100). Payload variants are reported in **Supplementary Table S14** and QC calls are summarized in **Supplementary Table S2**.

### mESC clone sequence verification

To verify correct genomic engineering of mESC clones, mean sequencing coverage was calculated for the genomic regions corresponding to the payload and non-payload (as defined above); and for the payload YAC/BAC backbone, landing pad (LP), LP backbone, and pCAG-iCre custom references. Terminal clipping of 400 bp was performed as described above for all genomic regions. Coverage was normalized by the mean genomic coverage of the engineered region flanks (regions captured by the *Sox2* bait, but unmodified) such that a single copy corresponds to a value of ∼0.5. For LP clones we expect normalized coverage of 0.5±0.25 for the non-payload region and <0.1 for LP backbone. For payload clones we expect normalized coverage of 1±0.25 for the payload region, 0.5±0.25 for the non-payload region and <0.1 for the LP, payload YAC/BAC backbone and pCAG-iCre. Human orthologous sequences, and regions smaller than 200 bp after clipping were ignored. Samples where the payload region was ignored were classified as “No call”. Coverage analysis results and QC calls for each clone are reported in **Supplementary Table S15** and summarized in **Table 1**.

We further verified the presence of BL6 allele variants, which are lost in regions replaced by landing pads and restored by delivered payloads, by calculating allelic ratios as the mean proportion of reads supporting the reference (BL6) allele (propREF) for the genomic regions described above. Values of 0.5±0.2 and 0±0.2 were expected for payload and non-payload regions, respectively. Regions with 10 or fewer variants were ignored. Samples where the payload region was ignored were classified as “No call”. Allelic ratio results and QC calls for each clone are reported in **Supplementary Table S15** and summarized in **Table 1**.

To verify genomic integration sites, we detected sequencing read pairs mapping to two different reference genomes using *bamintersect* as previously described^25^ with slight modifications: Samestrand reads mapping within 500 bp were clustered, a minimum width threshold of 125 bp was required for reporting, and junctions with few reads (<1 and 5 reads/10M reads sequenced for LP and payload samples, respectively) were excluded. *Bamintersect* junctions were classified hierarchically based on position (**Supplementary Table S16).** Results and QC calls for each clone are reported in **Supplementary Table S17** and summarized in **Table 1**.

### RNA Isolation, cDNA synthesis and mRNA expression analysis by real-time qRT-PCR

RNA was isolated from fresh or frozen cells using the Qiagen RNeasy-mini protocol. Since *Sox2* has no introns, additional steps were taken to ensure that RNA was not contaminated with trace residual genomic DNA. DNase treatment was performed on extracted RNA with the Turbo DNA-free kit (Ambion, Fisher AM1907) using the “rigorous DNase treatment” protocol prescribed by the manufacturer. Following DNase treatment, cDNA was synthesized from 1-2 µg total RNA with the Multiscribe High-Capacity cDNA Reverse Transcription Kit (Fisher 4368814), including a “-RT” no-reverse-transcriptase control for a subset of samples.

Real-time quantitative reverse transcription PCR (qRT-PCR) was performed using KAPA SYBR FAST (Kapa Biosystems KK4610) on a 384-well LightCycler 480 Real-Time PCR System (Roche) and threshold cycle (Ct, also called Cp) values were calculated using Abs Quant/2nd Derivative Max analysis. Primers were designed to detect *Sox2* in an allele-specific manner (**Supplementary Table S18**). An Echo 550 liquid handler was used for loading as described above. Thermal cycling parameters were as follows: 3 min pre-incubation at 95 °C, followed by 40 amplification cycles of 3 sec at 95 °C, 20 sec at 57 °C and 20 sec at 72 °C. For a subset of samples, the -RT controls were verified for lack of amplification and all had Ct>31.

Assays were performed in duplicate and replicate wells on the same plate were averaged after masking wells with no or very low amplification. Raw Ct values are listed in **Data S2**. Ct values for the BL6 *Sox2* allele ranged between 20 (WT) and 30 (ΔSox2), suggesting an accurate quantification range of 2^10^. ΔCt values for the BL6 and CAST *Sox2* alleles were computed relative to *Gapdh*. Replicate ΔCt measurements of the same clone across different plates were averaged. *Sox2* fold change was defined as the difference between the BL6 and CAST *Sox2* alleles, calculated as 2^ΔCt^[CAST–BL6]. Fold change was scaled to yield expression values ranging from 0 (ΔSox2) to 1 (WT) by subtracting the mean fold change calculated for the ΔSox2 samples from all data points, then dividing by the mean fold change of the appropriate WT payload samples (full locus or LCR) (**Supplementary Table S4**).

### Modeling

We fitted linear regression models to predict *Sox2* expression based on DHS composition and configuration using R v3.5.2^59^. Models were compared based on Bayesian Information Criterion (BIC) to evaluate the performance of predictor combinations. The composition of proximal (DHSs 1-8 and DHSs 10-16) and core LCR (DHS23, DHS24, CTCF25, and DHS26) regulatory elements was represented by their copy number in the payload. Regulatory element configuration was represented by indicator variables of their relative order (DHS24-DHS23 and CTCF25-DHS24; read as DHS24 before DHS23 and CTCF25 before DHS24) and sequence inversion (inv_DHS23, inv_DHS24, inv_CTCF25, and inv_DHS26). Payloads which did not map clearly onto these features (*Δ26-27 & 28, Δ20-23.3*, and the synthetic payloads in **Fig. 6** and **Supplementary Fig. S6**) were excluded from the analysis. Interaction between two elements was represented by indicator variables for the presence of both elements (DHS23and24, DHS23and26, and CTCF25andDHS26).

### Transcription factor motif analysis

We used motif matches previously derived from scanning the reference genomes using FIMO v4.10.244 with TF motifs as previously described^60^.

### Data Availability

DNase-seq data were obtained from https://www.encodeproject.org for ES_CJ7 (ENCLB163SYJ, DS13320) and H7 human ESC (ENCLB449ZZZ, DS11909)^7^. CTCF ChIP-seq^29^ (GSM2259905), ChIP-seq data^4, 27, 28^ (GSM560343, GSM560345, GSM560350, GSM687280, GSM687282, GSM687285, GSM845236, GSM845238, GSM1082340, GSM1082341, GSM1082342), and ChIP-NEXUS^30^ for Zic3 (GSM4087824), Pbx (GSM4087823), Esrrb (GSM4087822), Sox2 (GSM4072777), Oct4 (GSM4072776), Nanog (GSM4072778), and Klf4 (GSM4072779), PRO-seq^31^ (GSE130691), and STARR-seq^26^ (GSM4261634) were obtained from the GEO repository.

### Code Availability

The processing pipelines for Capture-seq, ChIP-seq, and DNase-seq data are available on Github at https://github.com/mauranolab/dnase. All code for analyses herein is available upon request.

## Acknowledgements

We thank Brendan Camellato, Florrie Zhu, Leslie Mitchell, Sudarshan Pinglay, Weimin Zhang, and Yu Zhao for help with yeast assembly. This work was partially funded by National Institutes of Health (NIH) grants RM1HG009491 (to J.D.B.) and R35GM119703 (to M.T.M.).

## Author Contributions

R.B., A.M.R., M.S.H., J.D.B. and M.T.M. designed experiments; R.B., M.S.H., G.E., R.O., and R.D.L. performed experiments; C.C., M.S.H., N.S., R.B., and G.E. assembled DNA payloads; M.S.H., H.J.A., E.H., G.E. and R.D.L. performed sequencing; A.M.R. and M.T.M. performed computational analyses; R.B., A.M.R., and M.T.M wrote the manuscript.

---

**Supplementary Fig. S1.**
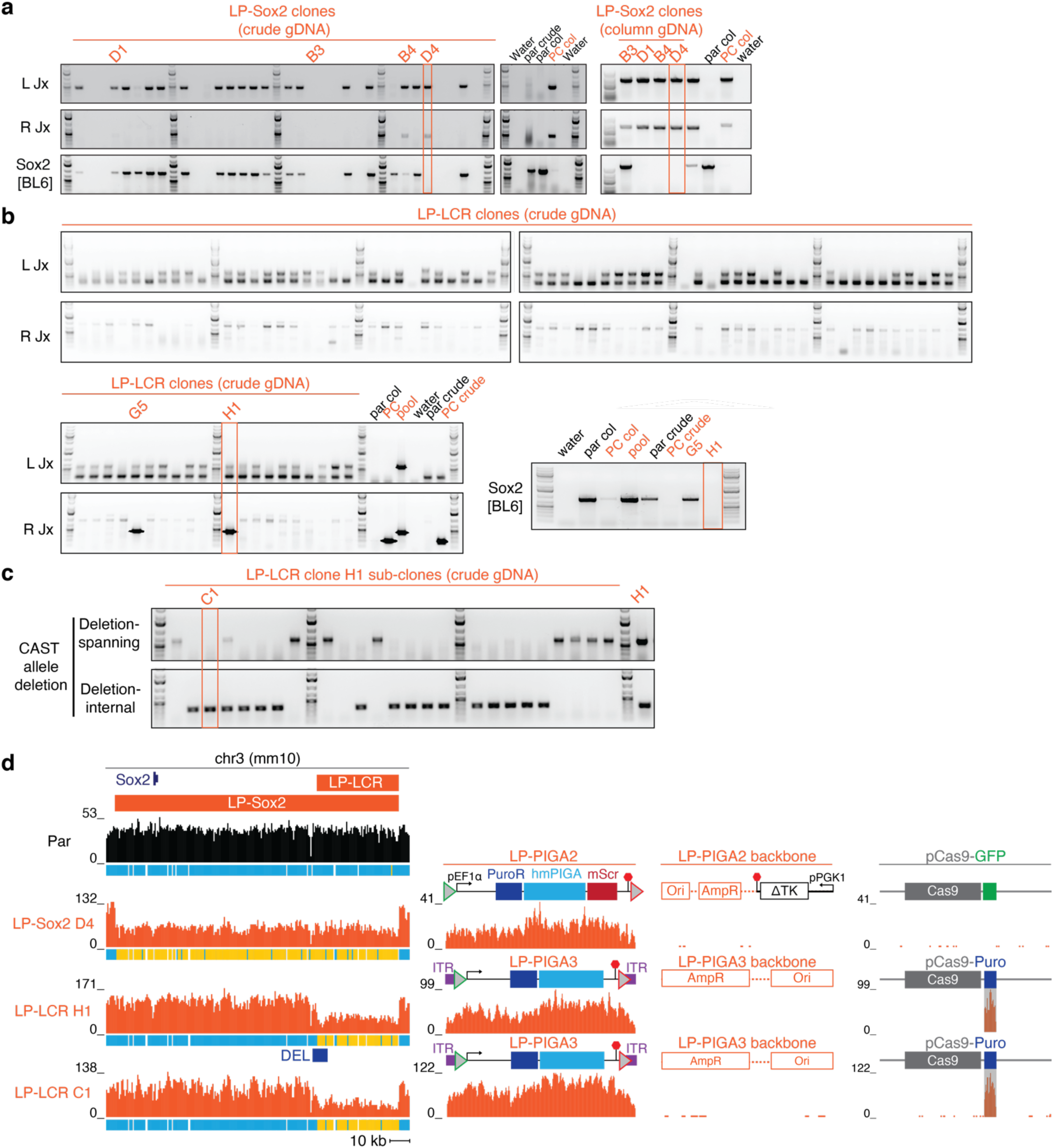
Landing pad integrations. **a**. Screening of BL6xCAST LP-Sox2 clones using PCR genotyping primers targeting the novel right junction (R Jx) and left junction (L Jx), as well as allele-specific primers to detect the loss of the targeted *Sox2* BL6 allele. Right, selected clones were revalidated using column gDNA. Clone D4 (orange rectangle) was selected for further verification by Capture-seq. Par, BL6xCAST Δ*Piga* mESCs; col, column gDNA; PC, BL6xCAST LP-Sox2 A1 mESCs^25^. **b**. Screening of BL6xCAST LP-LCR clones using PCR genotyping primers targeting the novel right junction (R Jx) and left junction (L Jx), as well as allele-specific primers to detect the loss of the targeted *Sox2* BL6 allele. Clone H1 (orange rectangle) was selected for further verification by Capture-seq. Par, BL6xCAST Δ*Piga* mESCs; col, column gDNA; PC, BL6xCAST LP-Sox2 A1 mESCs (which do not share the L Jx with LP-LCR clones and should produce a shorter amplicon from the R Jx). **c**. Screening of sub-clones of LP-LCR clone H1 using PCR genotyping primers spanning or internal to the CAST allele deletion. GeneRuler 1 kb Plus DNA Ladder (ThermoFisher Scientific) was used in panels **a-c**. **d.** Capture-seq analysis of parental Δ*Piga* mESCs, LP-Sox2 clone D4, and LP-LCR clones H1 and C1. Reads were mapped to the references indicated above. The PuroR gene in pCas9-Puro cross-maps with the LP-PIGA3 and is shaded gray. Ticks under each coverage track indicate heterozygous (blue) or homozygous non-reference (yellow) variants. ITR, inverted terminal repeats; pEF1α, human EEF1A1 promoter; PuroR, puromycin-resistance gene; hmPIGA, human mini PIGA gene; mScr, mScarlet; pPGK1, human PGK1 promoter.

**Supplementary Fig. S2.**
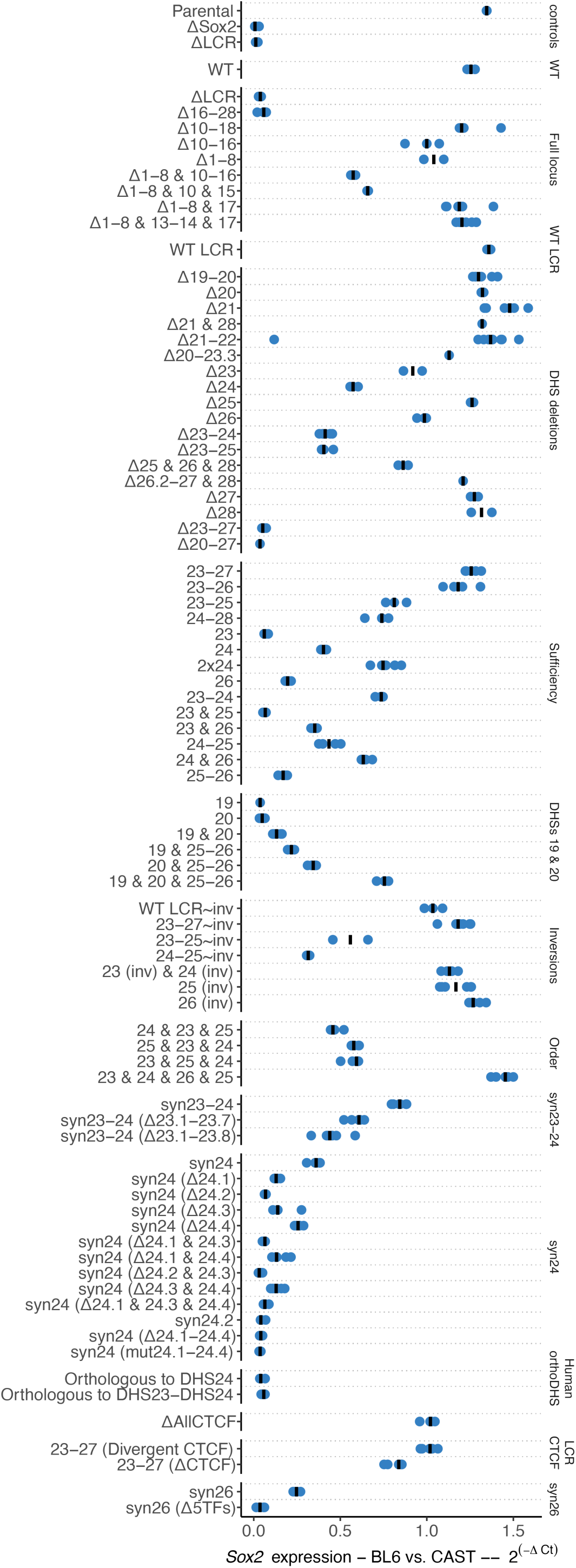
qRT-PCR results. *Sox2* expression from all mESC clones described in this manuscript. Displayed is the expression of the BL6 allele of *Sox2* relative to the CAST allele, calculated as 2^ΔCt[CAST-BL6]^. Points represent individual clones and bars indicate median.

**Supplementary Fig. S3.**
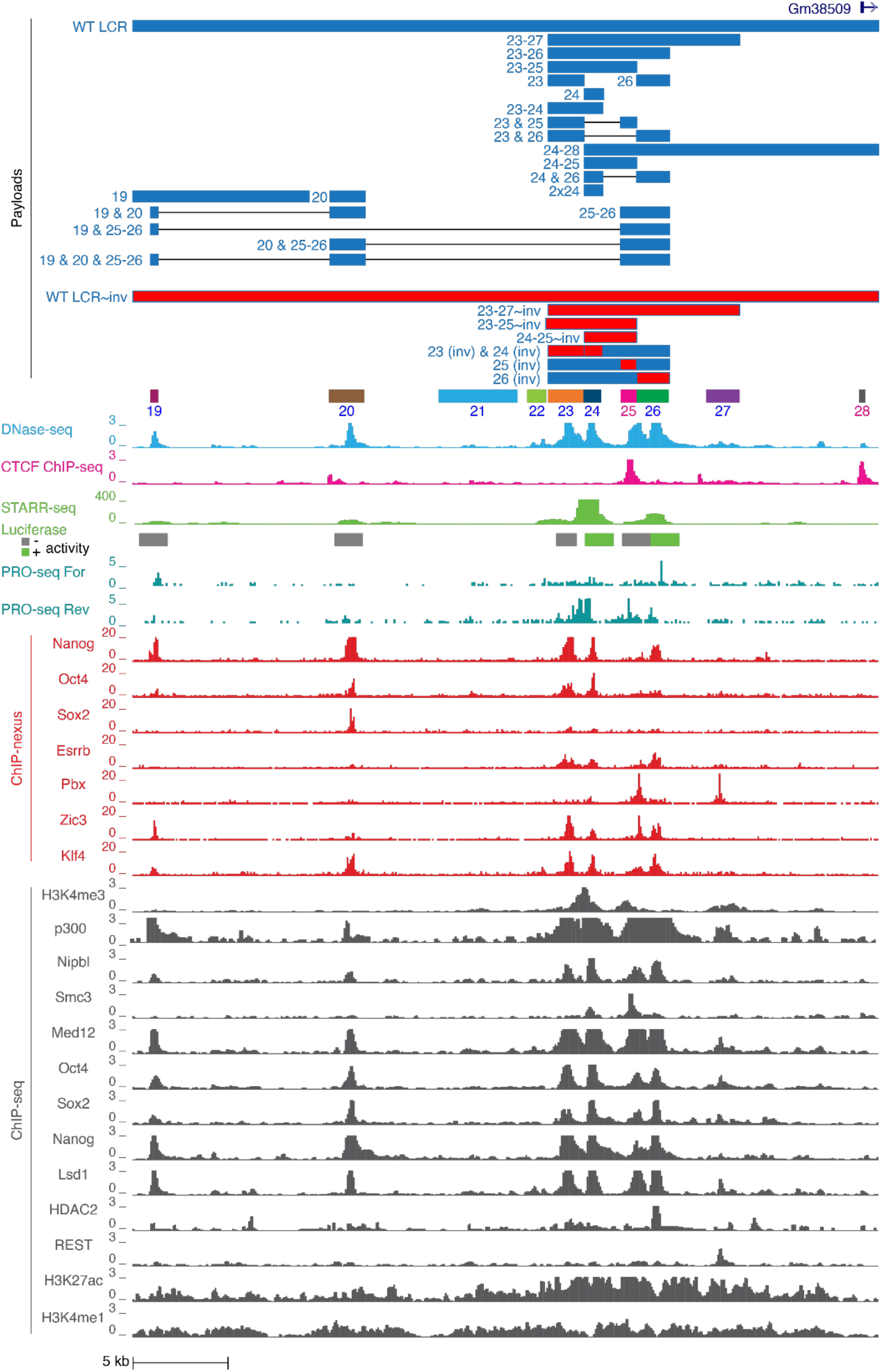
*Sox2* LCR structure and payloads. Browser shot for the *Sox2* LCR. DHSs and payloads included in **Fig. 3** are demarcated along with DNase-seq, ChIP-seq, STARR-seq, and ChIP-nexus tracks. See Data Availability for sources.

**Supplementary Fig. S4.**
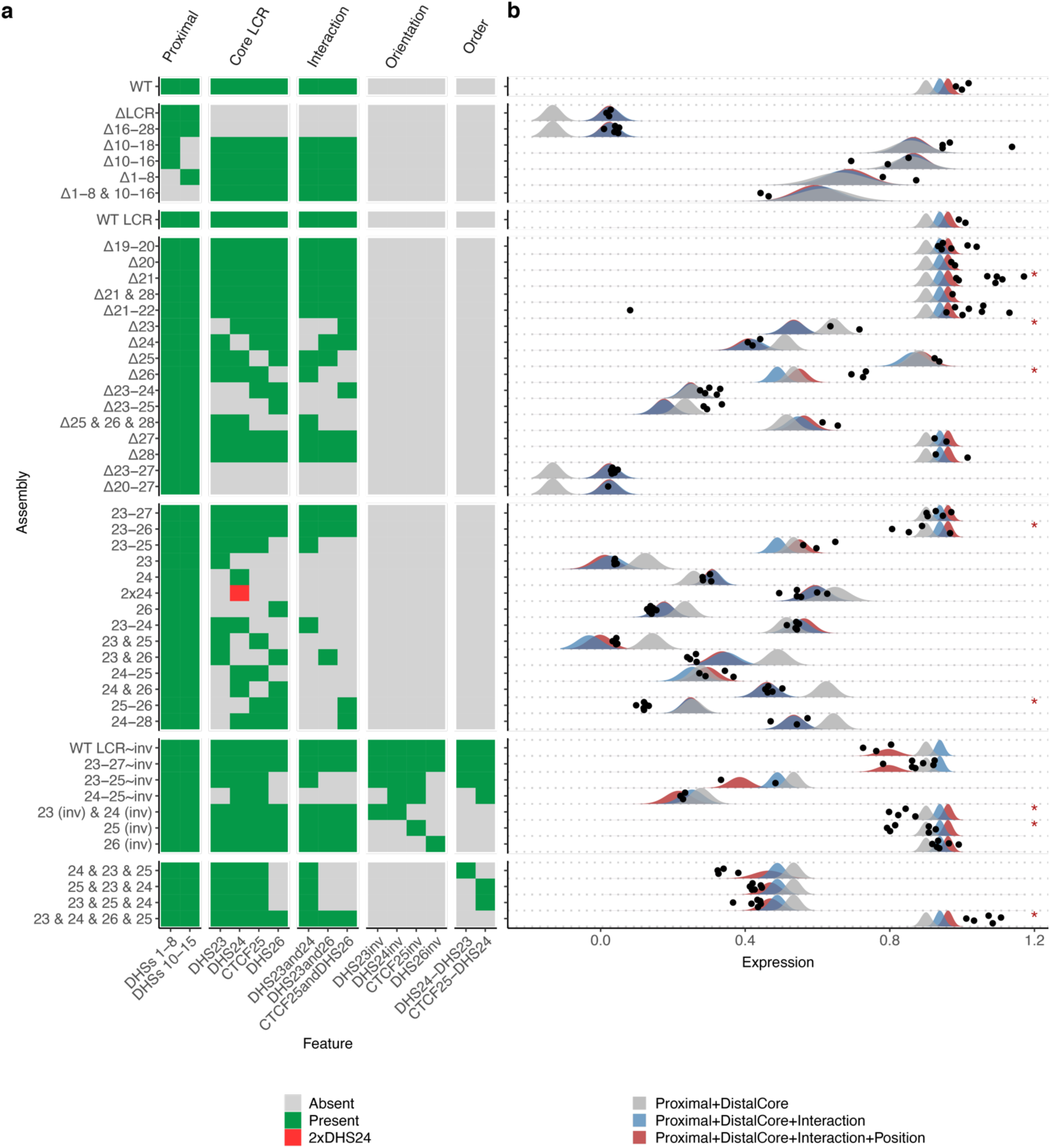
Linear regression model of *Sox2* regulatory architecture. **a.** Feature composition matrix. Gray, absence; blue, presence; red, two copies. **b.** Model predictions vs. actual data. Points represent the *Sox2* expression vs. WT for individual mESC clones. Distributions indicate the expected outcome distribution for each regression calculated based on Student’s t distribution using the estimated mean and standard error of each PL. Asterisks denote payloads for which measured data diverge from *Proximal*+*core LCR*+*Interaction*+*Position* model prediction by >5 standard errors.

**Supplementary Fig. S5.**
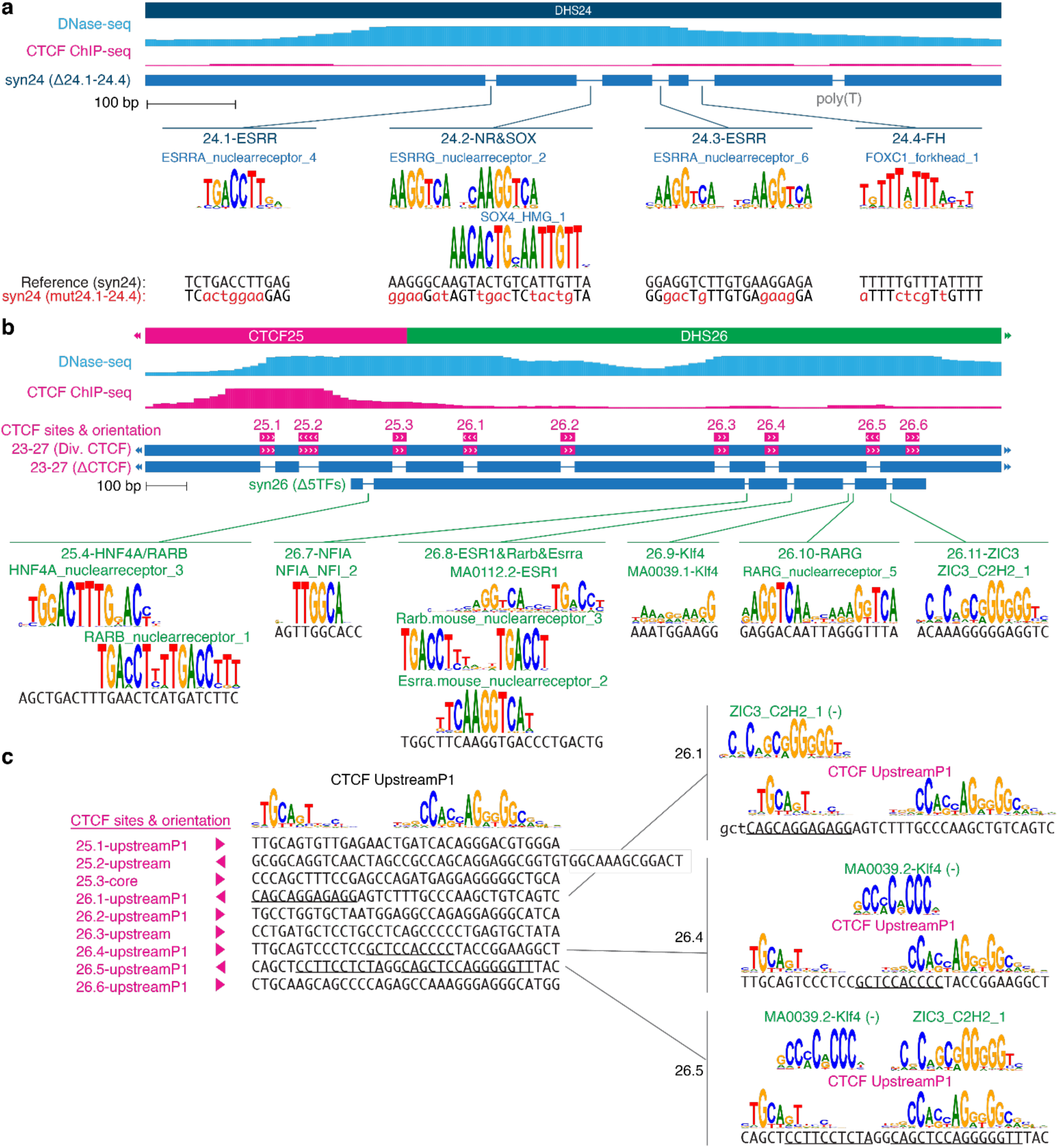
Schematic of TF-scale engineering. Schematic of TF site deletions and mutations in DHS24 (**a**) and TF site deletions and CTCF site inversions in CTCF25-DHS26 (**b-c**). Shown in **a** and **b** are DNase-seq and CTCF ChIP-seq data in mESCs. Selected payloads are shown including TF and CTCF sites, weblogos for matching motifs, and the actual reference (black). Mutated sequence in payload *syn24(mut24.1-24.2)* is in red. **c**. Reference sequences for 9 CTCF sites are shown aligned to the CTCF motif. Orientation in the reference is indicated with triangles. CTCF extended binding modes are represented using three models including the core sequence, an additional upstream sequence, and the up-stream sequence at an additional 1-bp spacing (upstreamP1)^42^. Recognition sequences for other TFs that over-lap CTCF recognition sequences are underlined and shown in detail at right and reverse orientation is indicated as (-).

**Supplementary Fig. S6.**
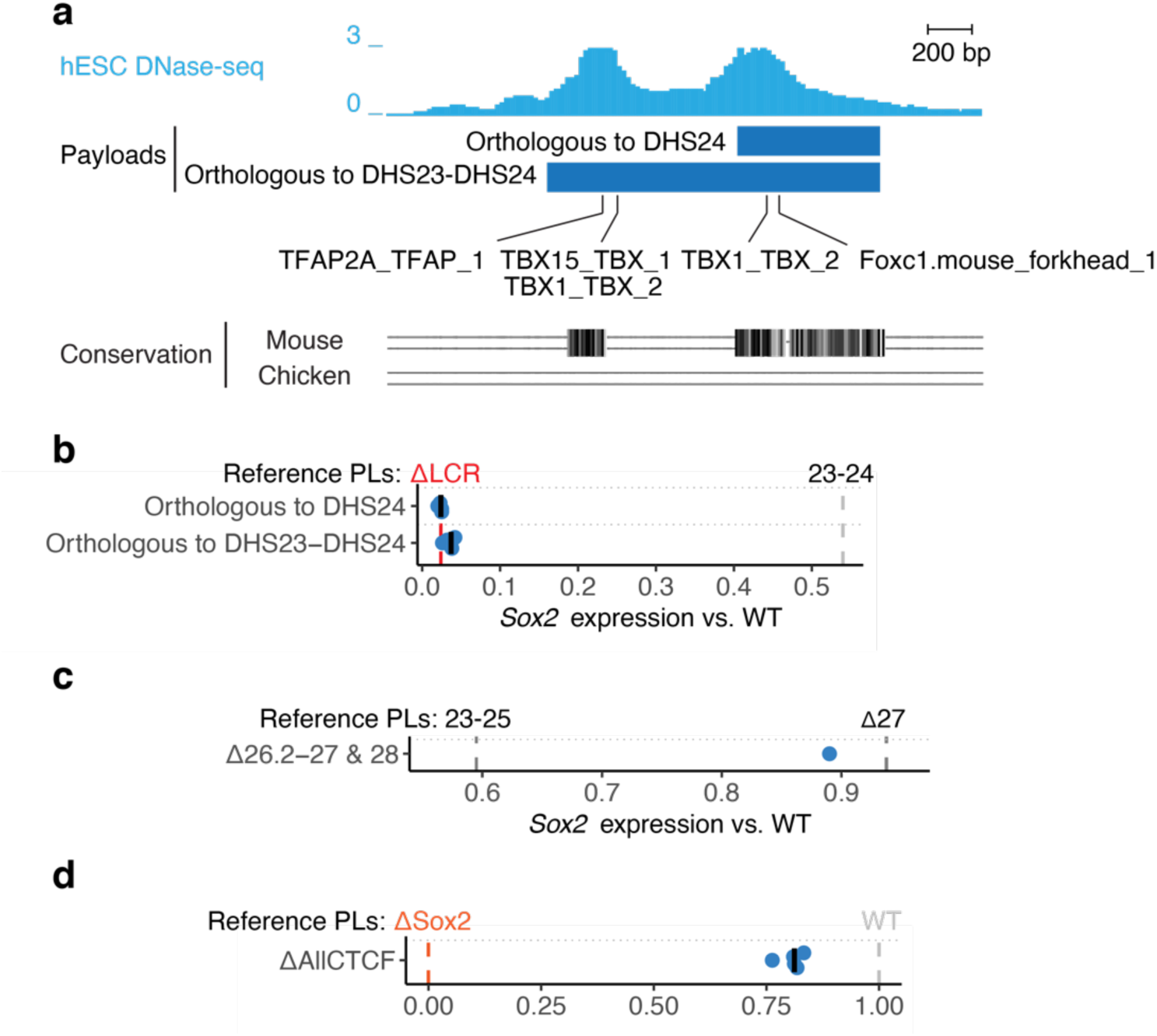
Core LCR and human orthologs analysis. **a.** Human DHSs orthologous to mouse *Sox2* DHS23 and DHS24. The mouse genomic coordinates of all *Sox2* regulatory regions were broken into 50-bp non-overlapping blocks. These blocks were remapped to the human genome (hg38) using UCSC liftOver with default options and adjacent remapped blocks within 200 bp were merged. Blocks were required to map reciprocally back to the original location in mouse using the same strategy. Shown are DNase-seq in H7 human ESC^7^, predicted TF recognition sequences, and phylogenetic conservation. Blue bars demarcate cloned human genomic regions. **b-d**. *Sox2* expression analysis. Each point represents the expression of the engineered *Sox2* allele in an independent mESC clone relative to baseline payloads (dashed vertical lines). Bars indicate median. Expression was scaled between 0 (*ΔSox2*) to 1 (*WT LCR*), and vertical gray lines indicate median expression of relevant baseline payloads. **b.** Activities of payloads comprising human *SOX2* regions orthologous to DHS24 and DHS23-DHS24 delivered to LP-LCR mESCs. **c.** Activity of payload *Δ26.2-7 & 28* (shown in detail in **Fig. 6b**). Expression data is reproduced from Fig. 3 and presented here on a different scale in comparison to the activities of payloads *Δ23-25* and *Δ27*. **d.** Activity of payload *ΔAllCTCF* which includes deletion of DHSs 1-8, CTCF sites 13-14, 17, and CTCF recognition sequences 25.1-26.6.

## SUPPLEMENTARY TABLES

**Supplementary Table S1. Payloads and assembly details.**

Strategies and reagents used for payload YAC/BAC assembly, as well as sequencing data. See Methods for details of each assembly strategy, Supplementary **Table S5** for cloning primer sequences, Supplementary **Table S8** for gRNA sequences, Supplementary **Table S9** for synthetic fragments sequences and Supplementary **Table S4** for payload labels used in figures. Sequence-verified unintended payload structures observed in bacteria or in mESCs are indicated in the Assembly strategy field and were assigned n/a values in relevant fields. For two payloads with detected variants (see Supplementary **Table S14**), two different YAC/BAC clones were delivered to mESCs.

This Table is provided as a separate file.

**Supplementary Table S2.** Summary of YAC/BAC QC. Summary Table for YAC/BAC QC analyses. See **Supplementary** Table S13 and **Supplementary** Table S14 for details.

|  | Assembled payload YAC/BACs (n=78) |  |  |
| --- | --- | --- | --- |
|  | <u>Passed</u> | <u>Failed</u> | <u>No call</u> |
| Sequencing coverage QC | 58 (74.4%) | 0 | 20 (25.6%)(<sup>a</sup>) |
| Sequencing variants QC | 78 (100%) | 0 | 0 |

<sup>a</sup> Small regions were not analyzed (see **Methods** for details).

**Supplementary Table S3. mESC clones.**

Summary Table for mESC clones described in this manuscript. Genetic Modification details DNA elements integrated and their integration site in square brackets; transiently transfected plasmids (e.g. pSpCas9 or pCAG-Cre) are indicated by null. See **Supplementary Table S4** for payload labels used in figures. “LP-PIGA A1” corresponds to “Bl6/C LP140(Sox2_1g_5g)_A1”, “LP-Sox2 D4” to “BL6/C LP300(Sox2_1g_5g)_201007_D4”, “LP-LCR H1” to “Bl6/C LP305(Sox2_6g_7g)_20200915_H1”, and “LP-LCR C1” to “Bl6/C LP305(Sox2_6g_7g)_20210326_C01”.

This Table is provided as a separate file.

**Supplementary Table S4. *Sox2* expression values and activity.**

*Sox2* expression in engineered mESC clones. The columns Sox2 (CAST) and Sox2 (BL6) list qRT-PCR ΔCt values relative to *Gapdh* for each *Sox2* allele. Fold change indicates the BL6/CAST 2^ΔCt^ ratio. Activity indicates scaled fold change as detailed in **Methods**. Payload ID, a unique name for each payload; Label, name as in figure panels.

This Table is provided as a separate file.

**Supplementary Table S5.**
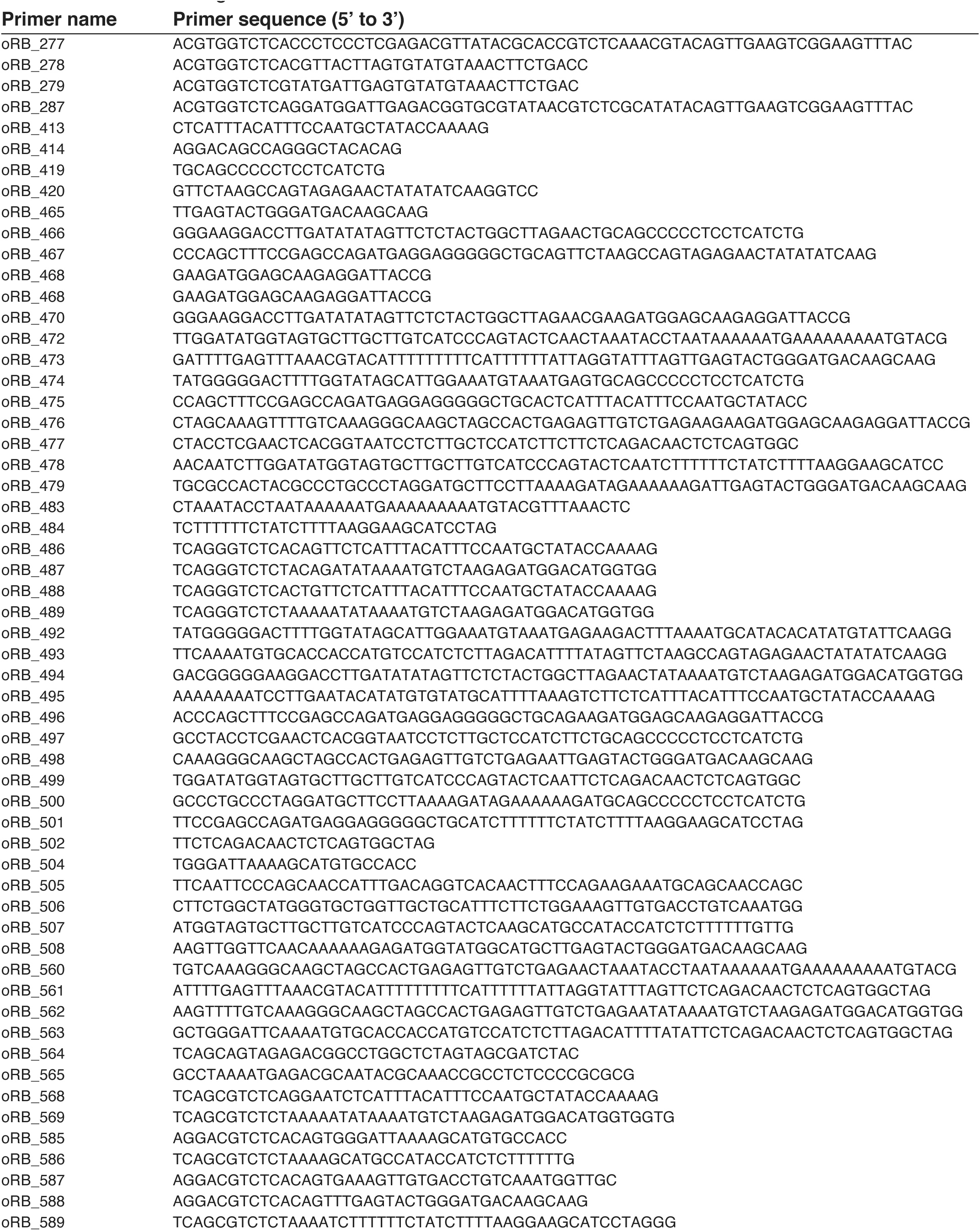
Cloning primers. Primers used for cloning.

**Supplementary Table S6. Genomic coordinates for engineered loci.**

Coordinates (mm10) for payloads, genomic deletions (marked with a “d”) and BACs. For some payloads, coordinates include mismatched regions derived from designed sequence mutation. See **Supplementary Table S4** for payload labels used in figures.

This Table is provided as a separate file.

**Supplementary Table S7.**
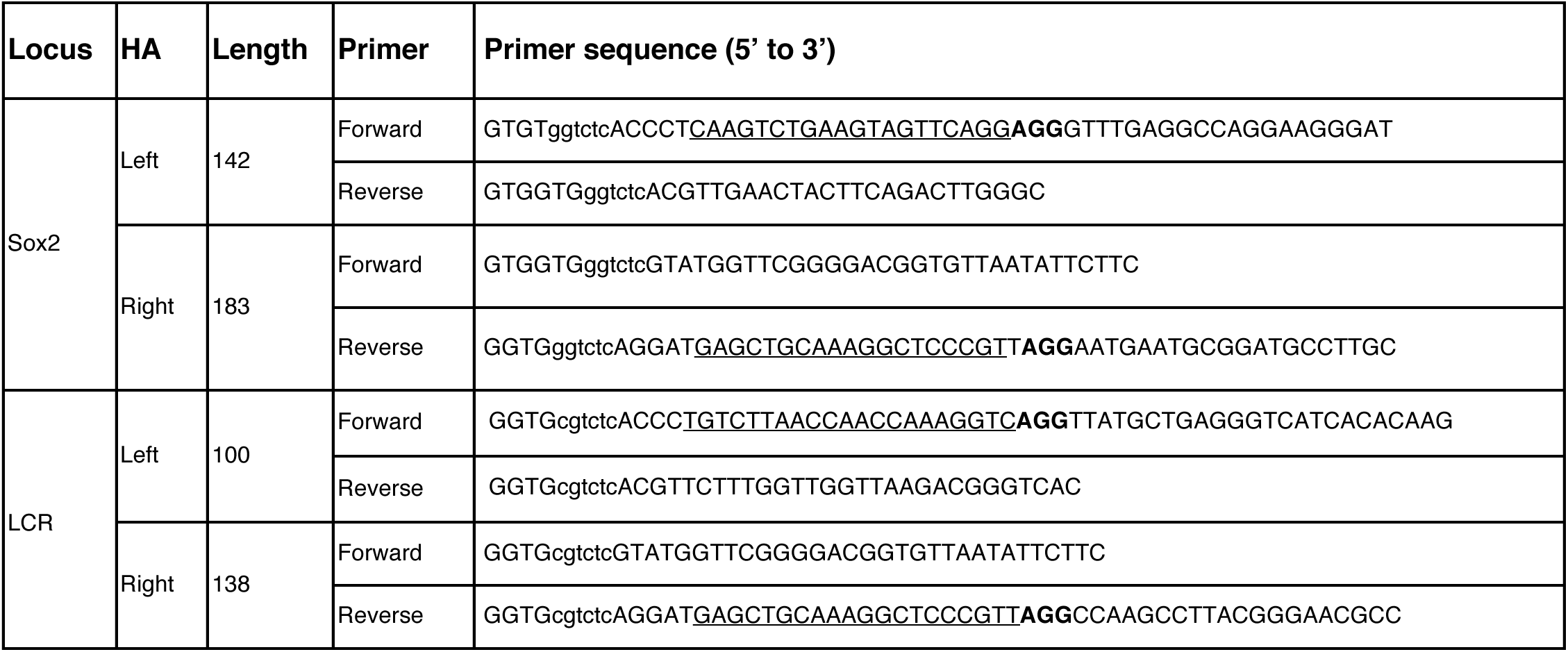
Homology arm cloning primers. Primer pairs used to clone left and right homology arms (HAs) using Golden Gate reactions Primer sequences contain BsaI or BsmBI sites in lower case, gRNA binding sites (underlined), and PAMs (bold). Length refers to the total size in bp of each HA.

**Supplementary Table S8.**
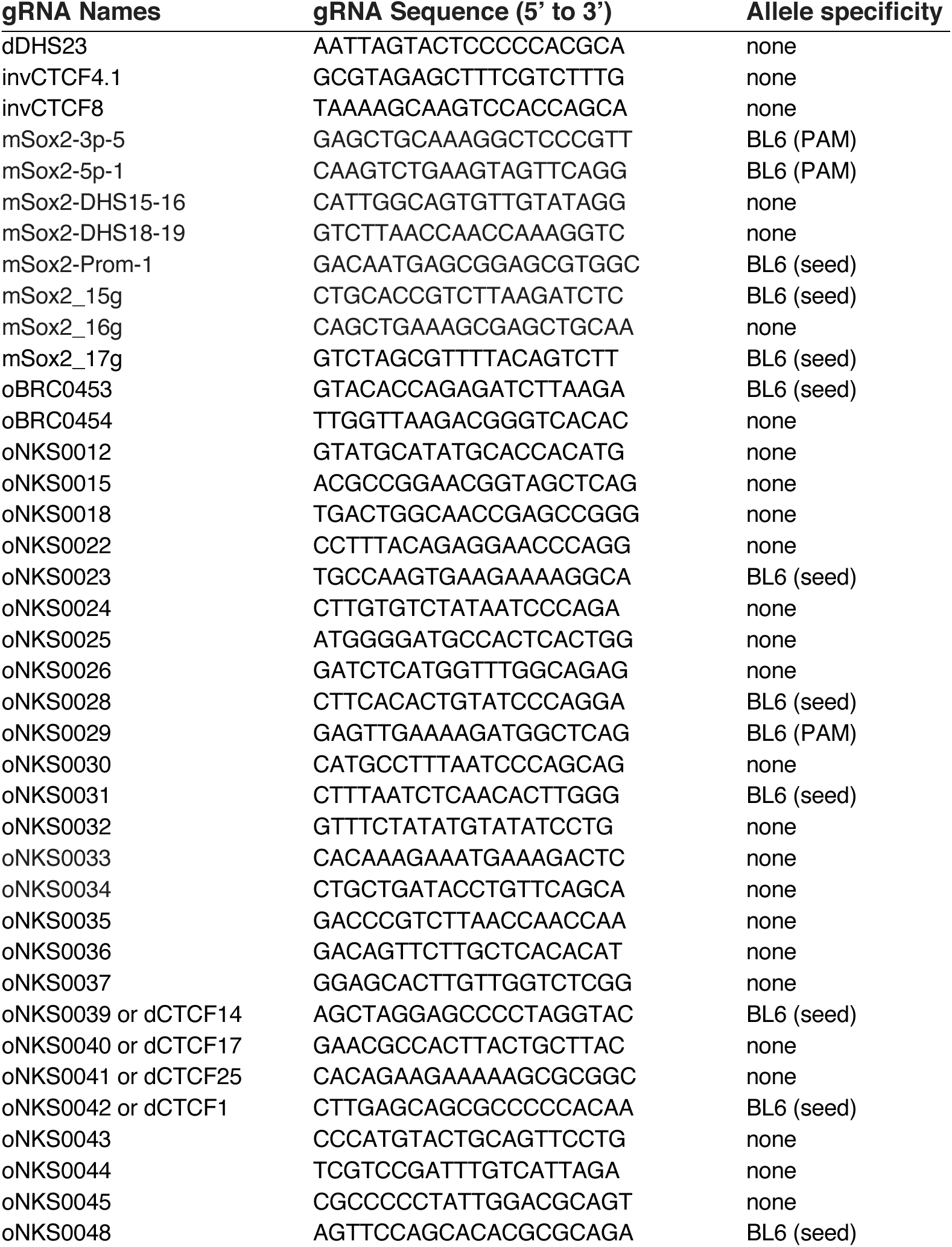
gRNAs. gRNAs used for LP integrations and payload assembly. Allele-specific gRNAs are identified as having CAST variants overlapping the PAM or gRNA seed sequence.

**Supplementary Table S9. Synthetic fragments.**

Sequences of synthetic fragments used for payload assembly.

This Table is provided as a separate file.

**Supplementary Table S10. LP and PL integration junction genotyping primers.**

Primer pairs used to verify newly formed junctions in engineered cells.

This Table is provided as a separate file.

**Supplementary Table S11.**
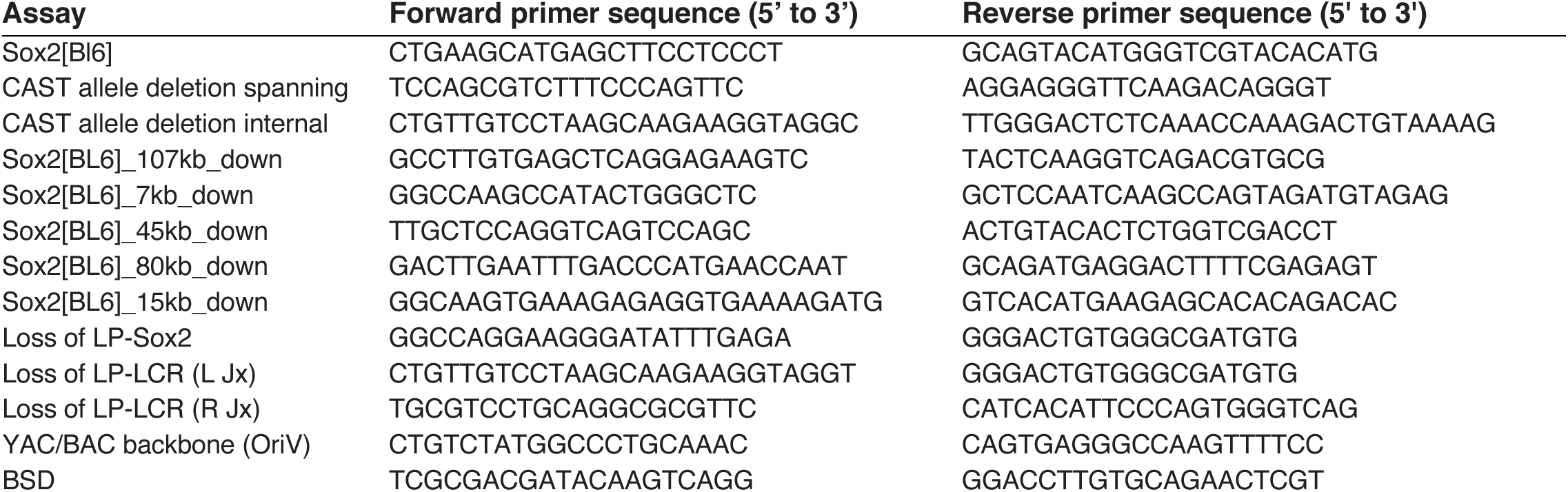
LP and PL integration additional genotyping primers. Primer pairs used to verify or characterize engineered cells.

**Supplementary Table S12. Sequencing libraries.**

Summary Table for DNA and DNA capture sequencing libraries. Sample ID, a unique sequencing library ID; Payload ID, a descriptor of the relevant landing pad or payload (see **Supplementary Table S4** for payload labels used in figures); Clone ID, a unique descriptor of mESC clones. For each sample, the number (#) of sequenced, analyzed and duplicate reads is provided. Bait Set is provided for capture samples.

This Table is provided as a separate file.

**Supplementary Table S13. YAC/BAC sequencing coverage QC.**

Summary Table for YAC/BAC payload DNA sequencing coverage depth analysis. Length (bp) and normalized (norm) coverage were calculated for regions corresponding to the payload and non-payload, as defined in **Methods**. Quality control (QC) calls are included with comments for unexpected values. See **Supplementary Table S4** for payload labels used in figures.

This Table is provided as a separate file.

**Supplementary Table S14. YAC/BAC variant calling.**

Summary Table for YAC/BAC payload variants relative to custom references. Ref, reference sequence; Alt, variant sequence; Qual, quality score; DP, sequencing depth; Feature, the custom reference feature overlapping the variant; RefSeq, variant and surrounding sequence; Comments, annotation and interpretation of variant location and relevance. None of the detected variants are expected to confound the measured activities of integrated payloads. See **Supplementary Table S4** for payload labels used in figures.

This Table is provided as a separate file.

**Supplementary Table S15. mESC clone capture sequencing QC.**

Summary of QC analyses of Capture-seq samples derived from engineered mESC clones, including sequencing coverage, allelic ratios, and integration site analysis (*bamintersect*). See **Methods** for more detail. Sample ID, a unique sequencing library identifier; Payload ID, LIMS name for relevant payload or LP; Clone ID, a unique descriptor of the mESC clone. Analyses were performed separately for: payload (genomic regions covered by the delivered payload), non-payload (genomic regions overlapping engineered locus and absent from payload), flanks (regions captured by the *Sox2* bait, but not engineered), PL[BB] (YAC/BAC backbone), LP (landing pad), LP[BB] (landing pad plasmid backbone), and iCre (pCAG-iCre plasmid). norm coverage, normalized mean coverage; propREF, proportion of reference to non-reference allele reads. QC calls for the coverage and variants analyses are included, as well as for the *bamintersect* analysis (see **Supplementary Table S16**). R/L Jx, right/left junction. See **Supplementary Table S4** for payload labels used in figures.

This Table is provided as a separate file.

**Supplementary Table S16.** *bamintersect* junction categories. Hierarchical criteria used to classify *bamintersect* junctions. bam1 and bam2 are the two references to which junctional reads were mapped. LP, Landing Pad; PL, payload; BB, backbone; Jx, Junction; Chr, chromosome.

| Clone Type | Priority | Jx class | bam1 |  |  | bam2 |  |  |
| --- | --- | --- | --- | --- | --- | --- | --- | --- |
|  |  |  | Chr | Position | Strand | Chr | Position | Strand |
| LP | 1 | L Jx | chr3 | 0-1000 bp upstream of HAL's 5' | + | LP | First 1000 bp | - |
| LP | 2 | R Jx | chr3 | 0-1000 bp downstream of HAR's 3' | - | LP | Last 1000 bp | + |
| LP | 3 | LP-BB | bam1 or bam2 maps to LP backbone |  |  |  |  |  |
| LP | 4 | off-target | None of the above |  |  |  |  |  |
| PL | 1 | L Jx<sup>(a)</sup> | chr3 | 0-1000 bp upstream of HAL 3' | + | PL | First 1000 bp | - |
| PL | 2 | R Jx<sup>(a)</sup> | chr3 | 0-1000 bp downstream of HAR's 5' | - | PL | Last 1000 bp | + |
| PL | 3 | L ITR-PL<sup>(b)</sup> | LP | First 1000 bp | + | PL | First 1000 bp | - |
| PL | 4 | R ITR-PL<sup>(b)</sup> | LP | Last 1000 bp | - | PL | Last 1000 bp | + |
| Any | 5 | Non-informative<sup>(c)</sup> | chr3 | Engineered region | Any | Any | Any | Any |
| Any | 6 | PL-BB<sup>(d)</sup> | bam1 or bam2 maps to YAC/BAC PL backbone |  |  |  |  |  |
| Any | 7 | off-target | None of the above |  |  |  |  |  |

<sup>a</sup> For payloads with wild type termini, CAST allele reads are indistinguishable from BL6 allele reads.
<sup>b</sup> Only applicable to payloads delivered to LP-PIGA3 (LP305) as the LP's ITRs remain flanking the integration site after PL delivery (see **Fig. 1**).
<sup>c</sup> Non-informative junctions often originate from the CAST allele within the engineered locus
<sup>d</sup> LP-BB and PL-BB junctions are often a result of contamination of gDNA samples by LP/PL plasmids.

**Supplementary Table S17. mESC *bamintersect* results.**

Summary of *bamintersect* integration site analysis for mESC clones capture sequencing samples. Each line describes a junction (Jx) detected by *bamintersect* (see **Methods**). Chrom (chromosome), chromStart, chromEnd, and Strand for bam1 and bam2 refer to the mapped reads location for the first and second reference genome, respectively. NearestGene indicates the nearest UCSC gene, including distance in bp (negative value indicates an upstream gene) for mm10 or nearest feature for custom references. ReadsPer10M is the number of read pairs supporting the junction per 10 million sequencing reads. Sample ID, a unique sequencing library identifier; Payload ID, LIMS name for relevant payload or LP; Clone ID, a unique descriptor of the mESC clone; Genomes indicate the reference genomes compared. Junc_type indicates the junction classification based on criteria detailed in **Supplementary Table S16**. See **Supplementary Table S4** for payload labels used in figures.

This Table is provided as a separate file.

**Supplementary Table S18.**
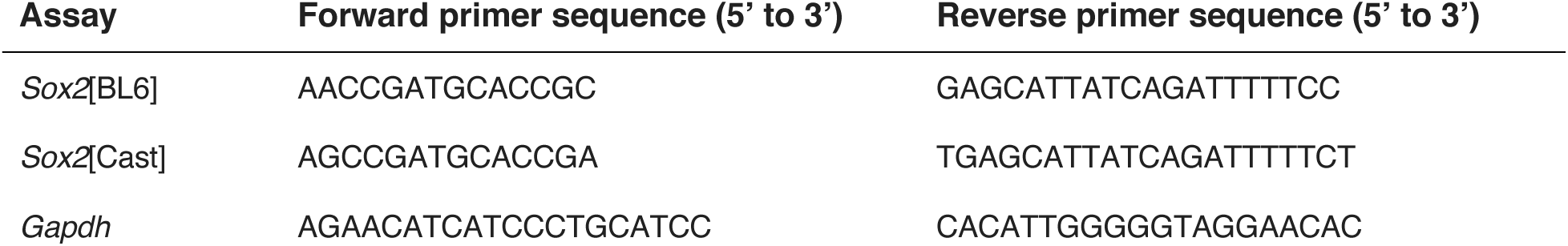
qRT-PCR primers. Primer pairs used for qRT-PCR.

## SUPPLEMENTARY DATA

**Data S1. Landing pad and payload sequences.**

Sequences for LPs and payloads described in this manuscript, provided in fasta format. For LPs, sequences include only the integrated region (excluding homology arms). For payloads, sequences include only the payload region of the YAC/BAC (from loxM to loxP) and correspond to the designed assembly (sequence variants are not included). See Supplementary **Table S4** for payload labels used in figures.

**Data S2. Raw qRT-PCR Ct values.**

Raw qRT-PCR Ct values for all mESC clones and assays. Source, the date when assay was performed; RT, indicates whether cDNA sample was prepared with a reverse transcriptase (TRUE) or without (“-RT” control, FALSE); Ct, threshold cycle. See **Supplementary Table S4** for payload labels used in figures.

